# Limited biological identifiability of dominance-labelled variance components in polyploid genomic prediction

**DOI:** 10.64898/2026.11.26.769574

**Authors:** Jinguang Tong, Qingchuan Yang, Mingna Li

## Abstract

Genomic prediction models for polyploid crops often partition genetic variance into additive and dominance-labelled components, but a fitted dominance component is not necessarily identifiable as digenic dominance (*D*_2_). We combined known-truth simulations with a panel-conditioned diagnostic workflow on 11 public datasets from alfalfa, potato, blueberry, guinea grass, rose, *Miscanthus*, sweetpotato and sugarcane. The workflow uses kernel geometry, permutation references, structure controls, planted-*D*_2_ spike-ins, posterior uncertainty and paired prediction tests. In autotetraploid simulations, the standard additive-plus-dominance model assigned appreciable variance to the dominance kernel when true *D*_2_ was absent, and its share responded only weakly to planted *D*_2_ signal. In crop panels, additive and dominance kernels were highly collinear, and Bayesian degree-kernel decompositions did not localize non-additive variance to *D*_2_, although restricted maximum likelihood placed it on *D*_2_ in alfalfa and potato. Observed shares exceeded matched permutation references by more than the pre-set margin in two of nine autotetraploid datasets, and no dataset met the criteria for biological *D*_2_ interpretation. A large fitted share was not sufficient for cross-validated prediction gain. We recommend reporting these components with diagnostics that separate biologically supported dominance, prediction-useful kernels and unresolved model-basis contributions.

**Plain Language Summary:** Many important crops, including alfalfa, potato and blueberry, carry four or more copies of each chromosome. Breeders use DNA markers to predict which plants will perform best, and the statistical models they use often report how much of the variation among plants comes from dominance, an interaction between gene copies. We tested whether that reported number can be trusted as a measure of real dominance. Using computer-simulated populations in which the truth was known, and eleven public datasets from eight crops, we found that the standard model reports a sizeable dominance share even when no dominance exists, and that real datasets rarely allow dominance to be told apart from other genetic effects. A dominance term can still improve predictions in some crops, but that is a separate question. We provide free software that checks, for any dataset, whether a reported dominance share can be interpreted biologically, is only useful for prediction, or cannot be resolved.

## 1 Introduction

Genomic prediction is increasingly used in autopolyploid and other dosage-based polyploid crops, including autotetraploid potato, alfalfa, blueberry, forage grasses such as *Panicum maximum*, and higher-ploidy crops such as sweetpotato and sugarcane (Batista et al., 2022; de Bem Oliveira et al., 2020; Endelman et al., 2018; Lara et al., 2019; Medina et al., 2025; Slater et al., 2016; Yadav et al., 2021). These crops often combine biological and practical features that make genomic prediction attractive: high heterozygosity, polysomic or partially polysomic inheritance, clonal or partially clonal propagation, expensive multi-year phenotyping, and long selection cycles. As allele-dosage genotyping has improved, genomic prediction models have moved beyond diploid-style marker coding and now routinely use dosage information to construct additive and non-additive genomic relationship matrices (Amadeu et al., 2023; Batista et al., 2022; Endelman et al., 2018; Slater et al., 2016). This has made it possible not only to predict breeding, clonal, or family performance, but also to partition fitted genomic variance into components labelled additive and dominance.

Biological non-additive variation matters for breeding, but the relevant signal is broader than digenic dominance alone. Classical theory partitions genetic variance into additive and non-additive components (Cockerham, 1954; Gallais, 2003; Kempthorne, 1957). A large additive component supports selection on transmissible breeding values and long-term population improvement, whereas substantial non-additive variation may motivate greater emphasis on clonal performance, hybrid or family performance, heterosis, mate allocation, or the maintenance of favourable heterozygous combinations. In crops where individual genotypes can be propagated directly, a non-additive component may also be useful for predicting clonal value even if it is not fully transmissible through gametes. Thus, for applied breeding, a dominance-labelled kernel can be useful whenever it captures genotype-similarity patterns that improve prediction, whether those patterns arise from classical dominance, heterozygosity, specific parental combinations, higher-order intralocus effects, or other sources of non-additive structure.

However, predictive usefulness and biological interpretation are different claims. The *D* component in a standard additive-plus-dominance *A* +*D* model is a fitted model component, not automatically an estimate of biological digenic dominance. Estimated genomic variance partitions depend on the parameterization of effects, family structure, the components included in the model, genotype-frequency assumptions, linkage disequilibrium (LD) and the standardization of relationship matrices (Lopes et al., 2015; Muñoz et al., 2014; Vitezica et al., 2013, 2017). Throughout this paper, we use “*D*-kernel share” to represent the fitted *D*-kernel fraction

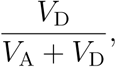

that is, the share of fitted genetic variance assigned to the *D* kernel among the fitted additive and *D* kernels. It is not a fraction of total phenotypic variance. We reserve “digenic dominance” for the degree-two intralocus contrast in the orthogonal framework, and use “*D* kernel” or “dominance-labelled kernel” for the fitted *D* component in a standard *A* + *D* genomic prediction model.

Autopolyploids make this distinction especially important. In a diploid biallelic locus there are three genotype states, so the classical additive–dominance decomposition has only one non-additive intralocus degree. In an autotetraploid biallelic locus, by contrast, allele dosage takes five values, 0, 1, 2, 3, 4, and a genotype-value function across these states can contain additive, digenic, trigenic, and quadrigenic components. More generally, ploidy *m* allows an intralocus hierarchy of dosage degrees from one to *m*. A standard dominance-labelled kernel therefore need not isolate biological digenic dominance in an autopolyploid. It may also absorb higher-order intralocus effects, heterozygosity-like patterns, or components correlated with additive relatedness. Orthogonal dosage contrasts provide a natural way to define additive and higher-order intralocus components, extending the logic of orthogonal quantitative-genetic decompositions and natural and orthogonal interactions (NOIA) parameterizations (Álvarez-Castro & Carlborg, 2007; Vitezica et al., 2017). For autotetraploids, an orthogonal-contrast model that dissects the genetic variance at a locus into independent com-ponents has already been developed (Chen et al., 2018); the contrasts used here extend that construction to any ploidy. But population-level orthogonality of contrast functions does not guarantee empirical separability of genome-wide kernels in finite, structured breeding panels.

The resulting gap is not the absence of dominance-aware genomic prediction models. Such models already exist and may improve prediction in some polyploid breeding settings (Amadeu et al., 2023; Batista et al., 2022; Endelman et al., 2018; Slater et al., 2016; Yadav et al., 2021). Nor is it new that a fitted dominance component need not improve prediction. In autotetraploid blueberry and potato, including digenic dominance explained part of the estimated genetic variance and improved goodness of fit, but predictive ability was similar to that of additive models (Amadeu et al., 2020). Song et al. (2024) built additive and dominance relationship matrices from allele dosage and found that, in blueberry, guinea grass (*Panicum maximum*) and sweetpotato, adding dominance did not raise prediction accuracy for most traits, which they attributed to dominance variance not being captured. Datasets from the same three public sources are among the panels analysed here.

This study therefore does not propose a new estimator or a new decomposition. It asks a question that the prediction comparisons leave open: when a model reports a dominance-labelled variance share, can that share be read as biological digenic dominance? The un-resolved question is whether the fitted dominance variance fractions reported from realistic autopolyploid panels are calibrated biological quantities. In particular, it remains unclear whether a large standard dominance share tracks true biological dominance, whether fitted non-additive variance can be localized to digenic rather than higher-order intralocus degrees, whether additive and dominance kernels are empirically separable in realistic breeding populations, and whether dominance-labelled variance components remain interpretable after accounting for population or family structure. It is also unclear whether a large reported dominance share predicts practical gain from adding a dominance kernel to a genomic prediction model.

Here we address these questions using ploidy-general orthogonal dosage contrasts and panel-conditioned diagnostics, with primary focus on autotetraploid panels. The framework constructs additive and higher-order intralocus contrast covariates and their corresponding genomic kernels, allowing the standard *A* + *D* model to be compared with an orthogonal *A* + *D*_2_ + *D*_3_ + *D*_4_ decomposition at ploidy four. We combine this framework with known-truth simulations, cross-crop reanalysis of public dosage-based polyploid datasets, kernel-collinearity diagnostics, effective-rank summaries, posterior uncertainty, phenotype-permutation nulls, structure-control analyses and cross-validated prediction comparisons. These analyses separate three related but distinct issues: population-level orthogonality of the contrast basis, sample-level separability of the fitted kernels, and finite-sample estimability of the variance components. The contribution is an integrated application of existing tools: known-truth calibration of the fitted share, and reference controls that are conditioned on each panel’s own genotype matrix.

Our central conclusion is not that dominance or non-additive biology is absent in autopolyploid crops. Rather, in the realistic panels examined here, the standard dominance share behaves primarily as a model-basis contribution unless additional diagnostics support biological interpretation. A dominance-labelled kernel may still be useful for prediction, and should be retained when cross-validation demonstrates a material benefit in the target population. But its fitted variance fraction should not be interpreted as biological digenic dominance merely because it is large. We therefore propose a diagnostic reporting workflow for distinguishing biologically supported *D*_2_ signal, prediction-useful dominance-labelled kernels and unresolved model-basis contributions.

## 2 Materials and Methods

### 2.1 Data sources and genotype preprocessing

We analysed dominance-labelled variance components using known-truth simulations and public polyploid genotype–phenotype panels. The empirical set comprised autotetraploid breeding and diversity panels, a same-individual blueberry dosage-calling check, and higher-ploidy sweetpotato and sugarcane case studies. Before model fitting, genotype records were aligned to analysed phenotypes, markers were filtered for basic quality, remaining missing dosages were mean-imputed, and dosage covariates were centred and scaled before kernel construction. Panel identities, phenotype adjustment rules, dosage-calling exceptions, exact marker filters, trait scopes and post-filter sample and marker counts are reported in Supplemental Methods and Supplemental Table S1.

### 2.2 Dominance-labelled and orthogonal degree kernels

For a biallelic locus in ploidy *m*, genotype dosage is *d* ∈ {0*, …, m*}. We constructed ploidy-general orthogonal dosage contrasts under the binomial random-mating reference

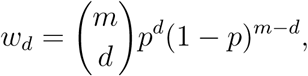

where *p* is the allele frequency. Marker-specific contrasts were obtained by Gram–Schmidt orthonormalization of 1*, d, d*^2^*, …, d^m^* under the weighted inner product

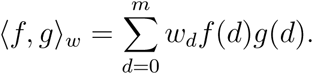

For marker *j*, *p* was estimated as 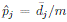, and the degree-*k* covariate for individual *i* was 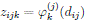. The first degree represents the additive dosage contrast; higher degrees represent intralocus non-additive dosage contrasts. In autotetraploids these correspond to additive, digenic, trigenic and quadrigenic contrasts. The degree-two contrast coincides with the classical digenic dominance deviation only under this reference, that is, random mating with random chromosome segregation and no double reduction, at the sample allele frequency. In panels that depart from the reference through inbreeding, selection, population structure or double reduction, *D*_2_ is a statistical degree-two dosage component defined relative to the reference, and the term is used in this sense for the empirical panels.

For each degree *k*, marker columns in *Z_k_* were centred to sample mean zero and scaled after imputation and filtering. The corresponding degree kernel was

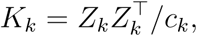

where *c_k_* rescales the kernel to mean diagonal one. The orthogonal model used *K*_1_*, …, K_m_*, with *K*_1_ denoted *K_A_* and, in autotetraploids, *K*_2_*, K*_3_*, K*_4_ denoted *K_D_*_2_ *, K_D_*_3_ *, K_D_*_4_.

The standard additive kernel was constructed from centred dosage covariates. The standard dominance-labelled kernel used the heterozygosity-style dosage function

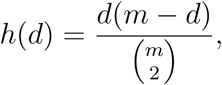

followed by the same marker centring, scaling and kernel normalization. At diploidy this is equivalent to the usual heterozygote indicator up to centring and scaling; at higher ploidy it is a dominance-labelled comparator kernel. Its fitted variance component was therefore interpreted as a standard *D*-kernel contribution, not as direct evidence of identifiable biological *D*_2_ dominance.

### 2.3 Simulation designs

Known-truth autotetraploid simulations were generated with AlphaSimR (Gaynor et al., 2021). A single coalescent founder pool with 450 founders and 2,400 single-nucleotide poly-morphism (SNP) markers was used to derive three *n* = 400 population structures: an un-related panel sampled from founders, a paternal half-sib panel with 50 families of eight individuals, and a full-sib panel with 50 families of eight individuals from random crosses among 40 founders. The panels shared marker and causal loci so that differences among them reflected relatedness structure rather than different genomic architectures. Meiosis used the AlphaSimR default of bivalent pairing, so double reduction was not simulated in these populations.

Traits were generated directly from the orthogonal contrast covariates. Pure additive, digenic, trigenic and quadrigenic traits placed effects only on the corresponding degree. A directional heterozygosity scenario used a genome-wide heterozygosity gradient and was not expected to localize to a single orthogonal degree. Genetic values were standardized within panel, and residual noise was added to target broad-sense heritability *H*^2^ = 0.6. The primary pure-architecture simulations used 50 replicates per architecture and population structure.

Mixed additive-plus-digenic simulations were used to test calibration. In each replicate, standardized additive and digenic genetic values, 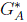 and 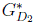, were generated from degree-one and degree-two contrasts. The genetic value was

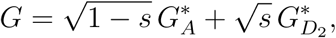

with *s* ∈ {0, 0.05, 0.10, 0.25, 0.50, 0.75}, followed by residual noise to target *H*^2^ = 0.6. Additive-only, standard *A* + *D* and orthogonal *A* + *D*_2_ + · · · + *D_m_*models were fitted per structure–fraction cell. Realized additive–digenic correlations were recorded and calibration was summarized by comparing fitted dominance fractions with the realized true digenic fraction.

A stress grid tested how calibration depends on sample size and marker number. Mixed additive-plus-digenic traits with *s* ∈ {0, 0.10, 0.25, 0.50} and *H*^2^ = 0.6 were simulated in full-sib panels with *n* ∈ {400, 1,200, 4,800, 19,200}, marker number *M* ∈ {1,000, 5,000, 20,000, 50,000, 100,000, 200,000} and two linkage-disequilibrium levels, with 50 replicates per cell. Implementation details of the simulations are provided in Supplemental Methods.

### 2.4 Empirical model fitting and prediction

For each empirical panel, additive-only, standard additive-plus-dominance and orthogonal degree-kernel models were fitted trait-wise. The general mixed model was

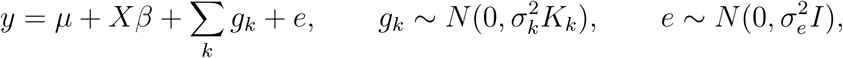

Here, *Xβ* denotes the fixed-effect term when population-structure covariates were included. The additive-only model used *K_A_*alone; the standard model used *K_A_*and the standard dominance-labelled kernel defined above; and the orthogonal model used the degree-specific orthogonal kernels *K*_1_*, …, K_m_*.

For the standard model, the dominance-labelled share was

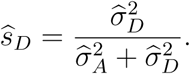

For the orthogonal model, fitted degree fractions were

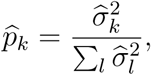

where the sum is over fitted genetic kernels. These fractions describe fitted genetic-kernel variance, not total phenotypic variance.

To assess dependence on the estimator, the first three traits of every panel were refitted under changed prior settings and by restricted maximum likelihood, with a profile-likelihood interval for the standard share (Supplemental Methods).

Predictive ability was assessed as the Pearson correlation between held-out phenotypes and predictions. Within each empirical panel, predictive ability was assessed by paired five-fold, five-repeat cross-validation with identical fold partitions across models. Held-out predictions were pooled over the five folds of each repeat and one correlation was computed per repeat; model differences were evaluated as paired contrasts of these repeat-level correlations. Phenotypes with missing values were analysed using the individuals available for each trait.

### 2.5 Diagnostic workflow for biological interpretation

We separated predictive utility from biological interpretation. A dominance-labelled kernel was considered predictively useful when it improved held-out prediction relative to the additive-only model in paired cross-validation. This criterion did not imply biological digenic dominance.

Biological interpretation was evaluated using panel-specific diagnostics. Kernel separability was measured by double-centred Frobenius cosines. For kernel *K*, let *K* = *HKH*, where *H* = *I* − **11**^T^*/n*. The cosine between two kernels was

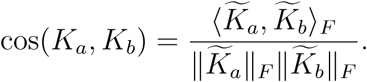

Values near one indicate that two kernels represent nearly the same empirical relationship direction. For a set of *q* kernels, we formed the *q* ×*q* cosine matrix *G*, with *G_ij_* = cos(*K_i_, K_j_*). If *λ*_1_*, …, λ_q_* are the eigenvalues of *G*, we report the effective rank

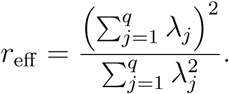

This quantity lies between 1 and *q*; larger values indicate more independent fitted kernel directions.

A phenotype-permutation null was used as a panel-level calibration reference. For each empirical panel, each selected analysed trait was permuted 30 times, the standard *A* + *D* model was refit, and the fitted dominance-labelled share was recorded. The permuted traits were the first three analysed traits of each panel. The observed share entering the comparison was computed on the same traits with the same point estimator and chain length as the permutations, and trait-level observed and null summaries were then averaged for panel-level workflow labels. This null preserves the genotype matrix and kernel geometry while removing genotype–phenotype association. It was used as a descriptive no-signal baseline rather than as a formal trait-level hypothesis test. Because permutation removes additive as well as non-additive signal, exceeding this null was treated only as a minimal condition and not as evidence for dominance. The reference that retains additive signal is the additive-only control: phenotypes were simulated on each panel’s own genotype matrix from additive effects alone (*H*^2^ = 0.6, 30 replicates), and real-trait *D*_2_ localization was judged against the distribution of this control. With 30 permutations per trait the null was used through its mean, whose standard error was about 0.006 or less per trait; tail percentiles of the null are too coarse to support a test and were not used in the labels. The location of this null near one-half is a property of the Bayesian estimator; under REML the share of a permuted phenotype usually falls at zero or one (96% of permutations; Supplemental Section C.3).

Structure-control analyses refit the standard model with leading genomic principal components (PCs) as fixed covariates. The control used the top ten PCs from the additive genomic relationship matrix. Where metadata were available, family, cohort or group kernels were also included and normalized to mean diagonal one. Structure sensitivity was summarized as

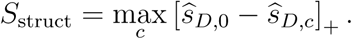

where 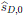 is the uncontrolled standard *D*-kernel share and 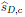 is the share under structure-control setting *c*.

The diagnostic workflow assigned fitted dominance-labelled variance to one of four reporting categories using optional, panel-conditioned reporting criteria. First, estimates were classified as structure-sensitive when the reduction in the fitted standard *D*-kernel share after structure control exceeded the structure-sensitivity threshold, either by an absolute margin or relative to the panel-specific structure-spike ruler. Estimates that survived this step were classified as *D*_2_-supported only when all biological-identification criteria were met: excess over the panel-specific phenotype-permutation null, recoverability of a planted *D*_2_ spike-in, real-trait localization toward the planted *D*_2_ direction rather than the additive-only null, and limited posterior uncertainty in the orthogonal *D*_2_ fraction. Estimates failing these biological-identification criteria were classified as *prediction-useful* when paired cross-validation showed a material and statistically supported gain from adding the standard *D* kernel, and as *model-basis* otherwise. The numerical thresholds used for each decision rule are listed in Supplemental Table S7. The thresholds are conventions and not calibrated error rates; the sensitivity of the labels to a 20% change in each threshold is reported in Supplemental Section C.5.

## 3 Results

### 3.1 Known-truth simulations separate standard D-kernel mislabelling from degree recovery

We first used known-truth autotetraploid simulations to ask whether the standard additive-plus-dominance model reports a large fitted *D* component only when biological digenic dominance is present. Across unrelated, half-sib and full-sib panels, the standard *A* + *D* model assigned a substantial share of fitted genetic variance to the *D* kernel under every simulated architecture (Figure 1A). This occurred not only for purely digenic and heterozygosity-gradient traits, but also for purely additive, trigenic and quadrigenic traits, all of which contain zero true digenic dominance by construction.

**Figure 1:**
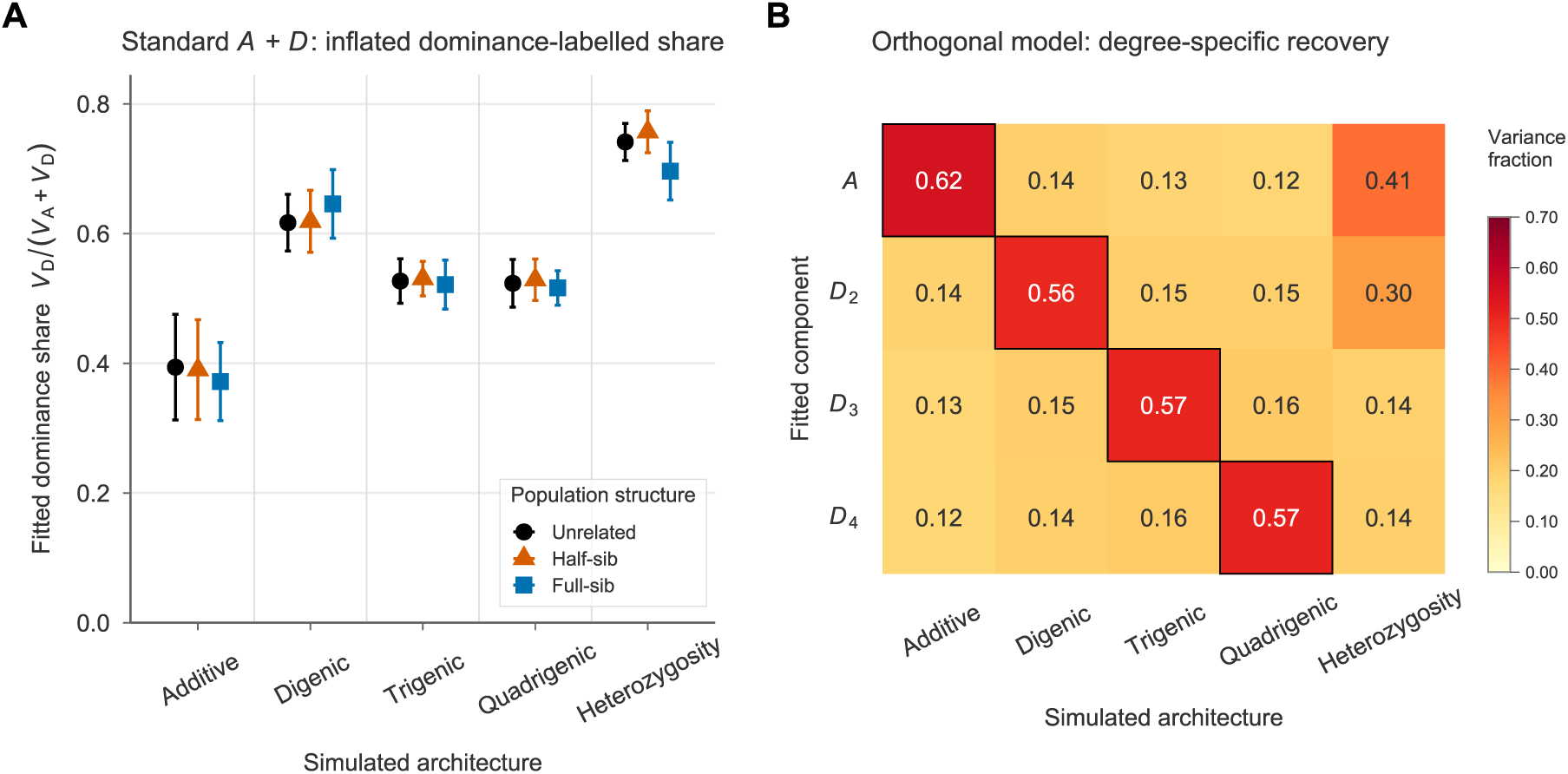
Known-truth simulations distinguish mislabelling from degree-specific recovery. **(A)** In autotetraploid simulations (*n* = 400, 50 replicates), the standard *A* + *D* model reports a large fitted dominance-labelled share, *V*_D_*/*(*V*_A_ + *V*_D_), even for additive, trigenic and quadrigenic architectures with zero true digenic dominance. Points show replicate means and error bars show 1 SD for unrelated, half-sib and full-sib population structures. **(B)** The orthogonal *A* + *D*_2_ + *D*_3_ + *D*_4_ model recovers the simulated dosage degree for single-degree truths, as shown by the boxed diagonal cells. Cell values are fitted variance fractions, normalized to sum to one across the four fitted genetic components (*A*, *D*_2_, *D*_3_, *D*_4_) within each simulated architecture, that is, down each column. Directional heterozygosity is not a single-degree architecture and loads mainly on additive and digenic components.

Thus, the fitted standard *D* component was not specific to true digenic dominance. Instead, it absorbed variance from additive effects, higher-order intralocus dominance and heterozygosity-like genome-wide gradients. A fitted share that remains large when true *D*_2_ is absent should therefore be interpreted as a dominance-labelled model component, not as a calibrated biological digenic-dominance fraction.

An orthogonal dosage-degree model *A* + *D*_2_ + · · · + *D_m_* behaved differently in the same controlled simulations. When the trait was generated from a single intralocus dosage degree, the largest fitted component matched the generating degree: additive traits loaded primarily on the additive kernel, digenic traits on the orthogonal *D*_2_ kernel and so forth (Figure 1B). Directional genome-wide heterozygosity was the exception. Because it represents a heterozygosity-gradient architecture rather than a single intralocus dosage degree, its fitted variance was distributed across more than one orthogonal component.

These simulations establish two points. First, the standard D-kernel share can be large even when true digenic dominance is absent, so it is not a specific estimator of biological digenic-dominance variance. Second, the orthogonal basis model can recover degree-specific signal when the data-generating architecture is aligned with the basis and the realized genotype panel provides sufficient separation among kernels. This positive-control result does not imply that the same components are separable or biologically identifiable in real breeding panels. That empirical question requires the panel-conditioned diagnostics analysed below.

### 3.2 Calibration curves show a high no-*D*_2_ baseline and shallow recovery

The pure-architecture simulations showed that the standard *A* + *D* model can assign a large share of fitted variance to the *D* kernel even when biological digenic dominance is absent. We next asked a stricter calibration question: as the known true *D*_2_ fraction increases from zero, does the fitted fraction increase in proportion to the truth? Using the same simulation framework, we generated mixed additive-plus-*D*_2_ traits as

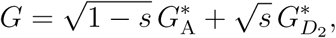

where 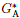 and 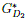 were standardized genetic values generated from the additive and or-thogonal *D*_2_ contrast bases, respectively. We then refit the standard *A* + *D* model and the orthogonal *A* + *D*_2_ + · · · + *D_m_*model. Under perfect calibration, the fitted fraction would equal the known true fraction on average, 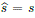; equivalently, a fitted-on-true calibration regression would have intercept zero and slope one (Figure 2).

**Figure 2:**
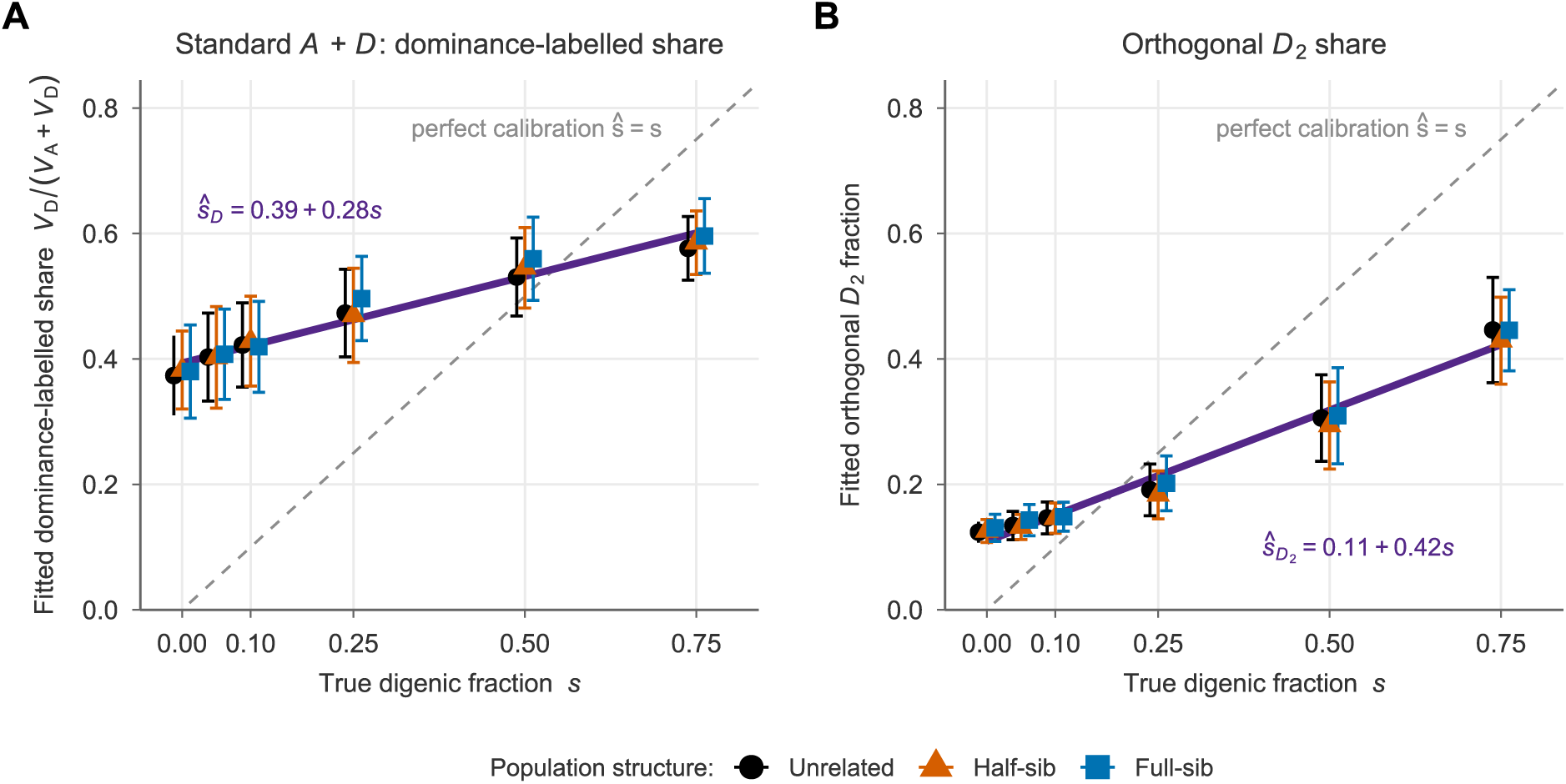
Calibration of fitted dominance fractions against the true digenic fraction. Mixed additive-plus-digenic autotetraploid traits were simulated with known true digenic fractions *s* (*n* = 400, *H*^2^ = 0.6, 100 replicates). Points show replicate means and error bars show 1 SD for unrelated, half-sib and full-sib panels. Dashed grey lines show perfect calibration, *s* = *s*; solid purple lines show pooled linear calibration fits across the three population structures. **(A)** The standard *A*+*D* model has a large no-dominance baseline: when *s* = 0, the fitted dominance-labelled share, *V*_D_*/*(*V*_A_ + *V*_D_), is already approximately 0.37–0.38, and the fitted relationship has a shallow slope, *s_D_* = 0.39 + 0.28*s*. **(B)** The orthogonal *A* + *D*_2_ + *D*_3_ + *D*_4_ model reduces the no-dominance baseline for the fitted *D*_2_ fraction, but the fitted relationship still falls below perfect calibration at high true digenic fractions.

The standard D-kernel share was not calibrated. Across the three population structures, the pooled calibration fit was

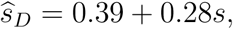

with unrelated, half-sib and full-sib panels showing the same qualitative pattern (Figure 2A). The positive intercept shows that the standard model assigned substantial variance to the *D* kernel even when no true *D*_2_ signal was present, and the shallow slope shows that additional true *D*_2_ signal was only weakly reflected in the fitted share. Thus, the standard D-kernel share is not a calibrated estimator of biological digenic-dominance variance; it is a fitted kernel contribution whose magnitude is only weakly related to the true *D*_2_ signal.

The orthogonal *D*_2_ fraction was better calibrated, but recovery remained incomplete. Its pooled calibration fit was

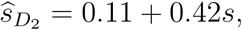

again with similar behaviour across population structures (Figure 2B). Isolating the orthogonal *D*_2_ contrast reduced much of the spurious baseline carried by the standard *D* kernel. However, the slope remained well below one, so increases in the true *D*_2_ fraction were still only partially recovered by the fitted *D*_2_ component. Orthogonalization therefore improved calibration, but did not by itself guarantee full recovery of true *D*_2_ variance in finite panels.

Residual under-calibration depended systematically on sample size, marker number and the realized separability of the fitted kernel basis. In the stress grid, increasing sample size reduced the no-*D*_2_ baseline and generally moved the fitted-versus-true calibration slope to-ward one. By contrast, increasing marker number *M* at fixed sample size generally worsened calibration, increasing the no-*D*_2_ baseline and reducing the calibration slope (Figure 3A,B). The geometry diagnostics showed the same pattern: calibration error increased with the additive–digenic Frobenius cosine and decreased with the effective rank of the four-kernel basis (Figure 3C,D). Thus, better empirical separation among fitted kernel directions was associated with lower calibration error. Together, these results indicate that finite-panel calibration is governed by empirical kernel separability: when additive and digenic directions are highly collinear, fitted dominance-labelled shares can show a positive no-*D*_2_ baseline and only a weak response to true *D*_2_ signal.

**Figure 3:**
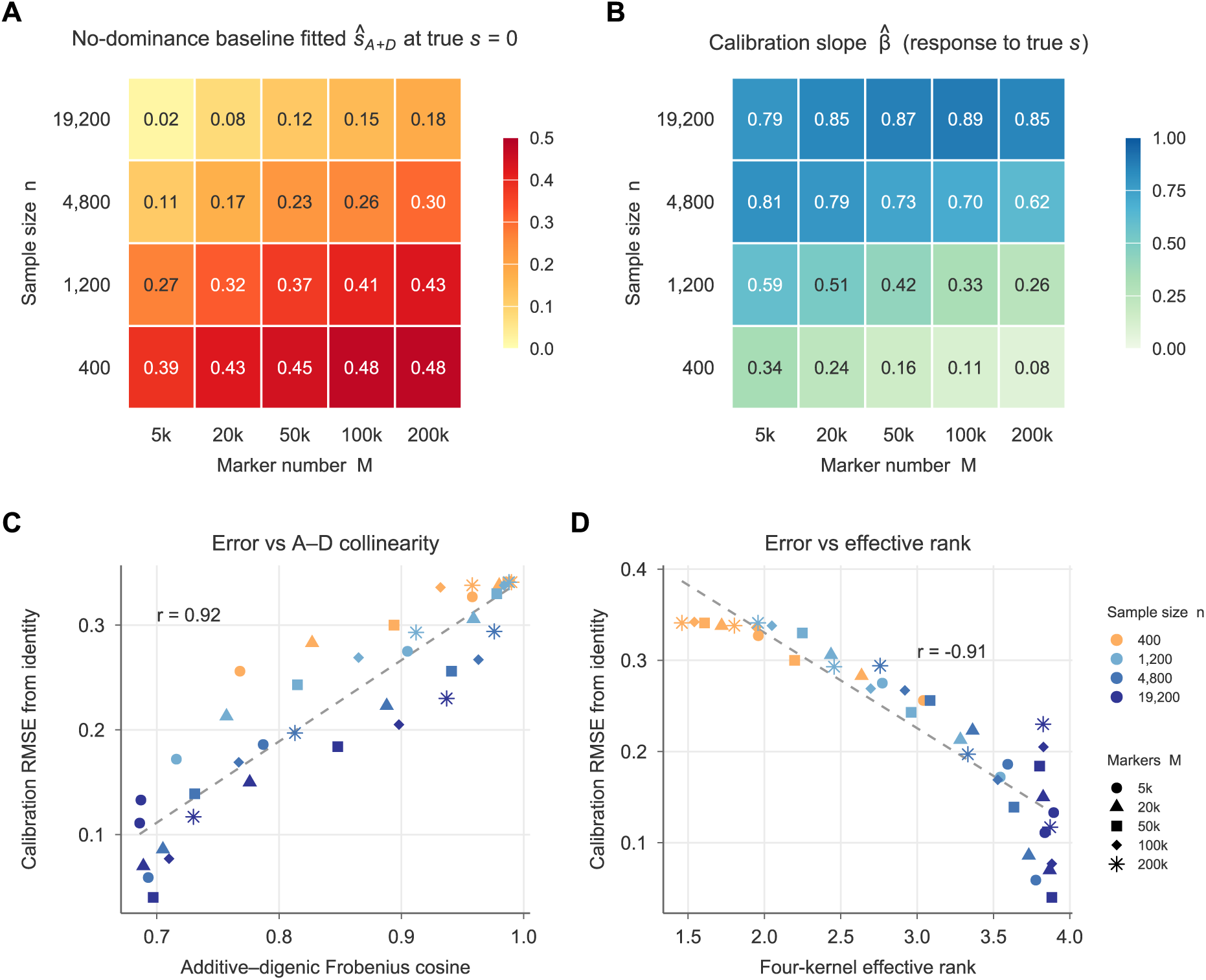
Finite-panel calibration improves with empirical kernel separability. All quantities are per-cell summaries over the full-sib stress grid (sample size *n*, marker number *M* 5,000, 20,000, 50,000, 100,000, 200,000, two linkage-disequilibrium levels, 50 replicates per cell; the grid also included *M* = 1,000, which is not shown). **(A)** No-dominance baseline—the fitted standard *A* + *D* dominance share at true digenic fraction *s* = 0—averaged over linkage disequilibrium across *n* and *M*; larger values are a larger spurious baseline. **(B)** Calibration slope *β*^^^, the response of the fitted share to the true fraction (regression of the fitted share on *s* 0, 0.1, 0.25, 0.5); values near one indicate a well-calibrated response. **(C,D)** Standard-share calibration root mean squared error (RMSE) from the *y* = *x* identity versus the additive–digenic Frobenius cosine (C) and the four-kernel effective rank (D); each point is one (*n, M,* LD) cell. The baseline grows and the slope degrades as the kernels become collinear (high cosine, low effective rank), driven by marker density relative to sample size.

### 3.3 Bayesian fits of autotetraploid panels do not localize non-additive variance to *D*_2_

We next asked whether the simulation-based warning signs also appear in empirical autotetraploid breeding and diversity datasets. We analysed eight independent public autotetraploid datasets plus one same-individual dosage-calling check, spanning alfalfa, blueberry, *Miscanthus*, *Panicum*, potato and rose (Figure 4). The blueberry UF-call dataset used the same individuals as the UF-GP dataset but an alternative dosage-calling pipeline, so we treated it as a robustness check rather than as an independent biological population. The full experiment results are reported in Supplemental Table S4.

**Figure 4:**
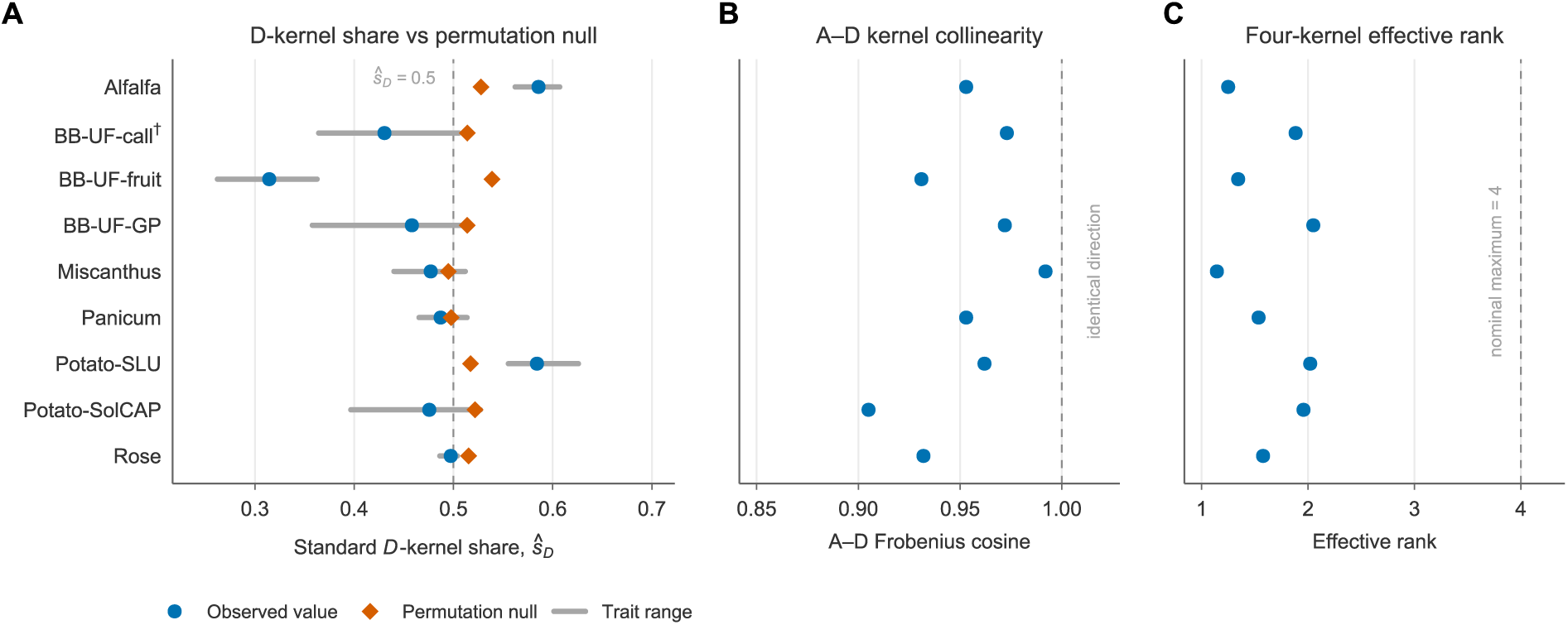
Autotetraploid empirical-panel diagnostics of fitted *D*-kernel shares and kernel separability. The analyses were applied to eight public autotetraploid datasets plus one dosage-calling check. Rows are aligned across panels. **(A)** Standard *D*-kernel share, *s_D_* = *V*_D_*/*(*V*_A_+ *V*_D_), from the standard *A* + *D* model compared with a phenotype-permutation no-signal null. Observed and null values are matched: blue circles show the observed share averaged over the traits used for the permutation null (the first three analysed traits of each dataset), estimated with the same point estimator and chain length as the permutations; grey horizontal lines span the minimum to maximum of those traits; and orange diamonds show the permutation-null mean for the same traits. The dashed vertical line marks 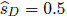. **(B)** Frobenius cosine between the additive kernel and the standard *D* kernel. Values near one indicate strong empirical collinearity between the two fitted kernel directions. **(C)** Effective rank of the four orthogonal *A* + *D*_2_ + *D*_3_ + *D*_4_ kernel set. All empirical datasets fall far below the nominal maximum of 4, indicating weak separability among orthogonal kernels.

The standard *A* + *D* model reported a substantial *D*-kernel share in every dataset, with trait-averaged values from the full-length fits (for alfalfa, its representative yield trait) ranging from 0.41 to 0.61 (Supplemental Table S4). The phenotype-permutation no-signal nulls were themselves centred near one-half, with dataset means from 0.49 to 0.54. We compared observed and null shares on the same traits and with the same estimator (Figure 4A; Supplemental Table S6). The observed share exceeded its null by more than the 0.05 margin used in the workflow in two datasets, alfalfa (+0.058) and potato SLU (+0.067). It was within 0.05 below the null in four datasets (*Panicum*, rose, *Miscanthus* and potato SolCAP) and further below it in the three blueberry analyses (−0.056 to −0.224). Because phenotype permutation preserves the realized genotype matrix while removing genotype–phenotype association, this null shows that a large fitted *D*-kernel share can arise from fitting the model to the realized dataset even without genetic signal. The relevant comparison is therefore excess over the dataset-specific no-signal null, not the absolute magnitude of the fitted *D*-kernel share.

Under the default-prior Bayesian model, the orthogonal decomposition did not resolve the non-additive degree components. Across datasets, posterior mean fractions assigned to *D*_2_, *D*_3_ and *D*_4_ were comparable rather than concentrated in *D*_2_. Full posterior summaries showed broad and mutually overlapping 95% credible intervals for the three non-additive degrees in every dataset (Supplemental Table S4). Thus, the fitted degree components were not empirically separable in these datasets. The result indicates poor localization of fitted non-additive variance to biological *D*_2_, rather than evidence for biological trigenic or quadrigenic dominance.

These fractions depend on the estimator. We refitted the first three traits of every panel with changed priors and with restricted maximum likelihood (REML), which uses no prior (Supplemental Table S5). Within the Bayesian model the prior settings mattered little: the standard share of a trait moved by at most 0.13, and the degree fractions stayed near-equal. REML differed in two ways. First, its 95% likelihood-ratio intervals for the standard share were wide. Of 33 empirical traits, 12 intervals covered the whole range from zero to one, and on permuted phenotypes the REML share fell at zero or one in 96% of permutations; the concentration of the Bayesian permutation shares near one-half is therefore a property of that estimator. Second, REML did not spread non-additive variance over the degrees. The REML fraction of degrees three and above was below 0.05 in 26 of 33 traits, and the REML *D*_2_ fraction was 0.49 to 0.55 for the three alfalfa traits and 0.19 to 0.33 for potato SLU, and at most 0.04 in *Panicum*, *Miscanthus*, rose, sweetpotato and sugarcane. The REML intervals excluded a zero standard share for ten traits, in alfalfa, both potato panels and blueberry UF-GP. The near-equal *D*_2_, *D*_3_ and *D*_4_ fractions are therefore a feature of the Bayesian fit, under every prior setting examined, and not evidence about degree composition. The REML degree fractions are point estimates without intervals, and REML localization was not calibrated against known truth, so they do not by themselves establish digenic dominance in alfalfa or potato.

Kernel geometry was consistent with this lack of localization. In every empirical dataset, the additive and standard *D* kernels were nearly collinear, as shown by high *A*–*D* Frobenius cosines (Figure 4B). The broader orthogonal *A*+*D*_2_+*D*_3_+*D*_4_ kernel set also had low empirical dimensionality, with effective ranks far below the nominal maximum of four (Figure 4C). Thus, the empirical genotype matrices provided limited information for separating additive, standard-*D* and orthogonal dosage-degree kernel directions.

Together, the permutation-null, orthogonal-decomposition and kernel-geometry results support the same interpretation: the standard fitted *D* component is a dominance-labelled model-basis contribution unless additional evidence demonstrates excess over a no-signal null, separability among fitted kernel directions and localization of non-additive variance to *D*_2_.

### 3.4 Predictive usefulness is separate from dominance-variance interpretation

Variance-component interpretation and predictive performance answer different questions. We therefore compared additive-only (*A*), standard *A* + *D* and orthogonal *A* + *D*_2_ +· · · + *D_m_*models using paired five-fold, five-repeat cross-validation across the empirical prediction datasets (Figure 5; per-dataset values in Table 1). The same fold partitions were used for all models within each dataset. Held-out predictions were pooled over the five folds of a repeat, one correlation was computed per repeat, and model differences were evaluated as paired contrasts of these correlations. Because trait numbers differed across datasets, we report paired prediction contrasts at dataset levels rather than using only a pooled mean that would be dominated by trait-rich datasets.

**Figure 5:**
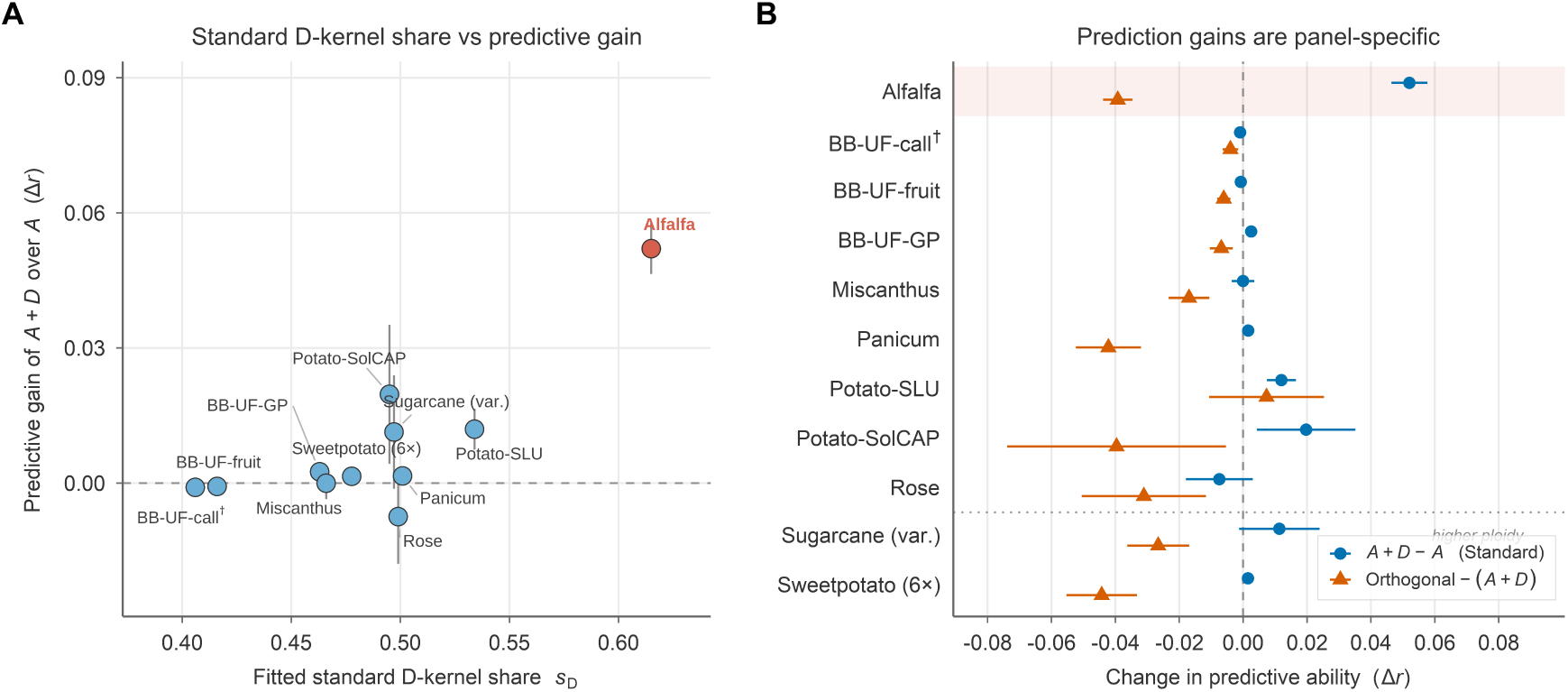
Predictive usefulness is separate from dominance-variance interpretation. Paired cross-validation compared additive-only (*A*), standard additive-plus-dominance (*A* + *D*) and orthogonal degree-kernel models within each panel. **(A)** Panel-level fitted dominance-labelled share, *V*_D_*/*(*V*_A_ + *V*_D_), plotted against the predictive gain of the standard *A* + *D* model over the additive model, Δ*r* = *r*(*A* + *D*) *r*(*A*). Panels have similar fitted dominance-labelled shares, but a substantial share did not guarantee predictive gain, which is near zero for most panels. **(B)** Paired changes in predictive ability by panel. Blue circles show the standard dominance-kernel contrast, (*A* + *D*) *A*, and orange triangles show the orthogonal-model contrast, Orthogonal (*A* + *D*). Horizontal intervals show 1 between-trait standard error of each panel-mean contrast. The shaded row highlights alfalfa, where the standard dominance kernel gave the clearest predictive gain. BB denotes blueberry; the dotted horizontal divider separates autotetraploid panels from higher-ploidy case studies.

**Table 1:**
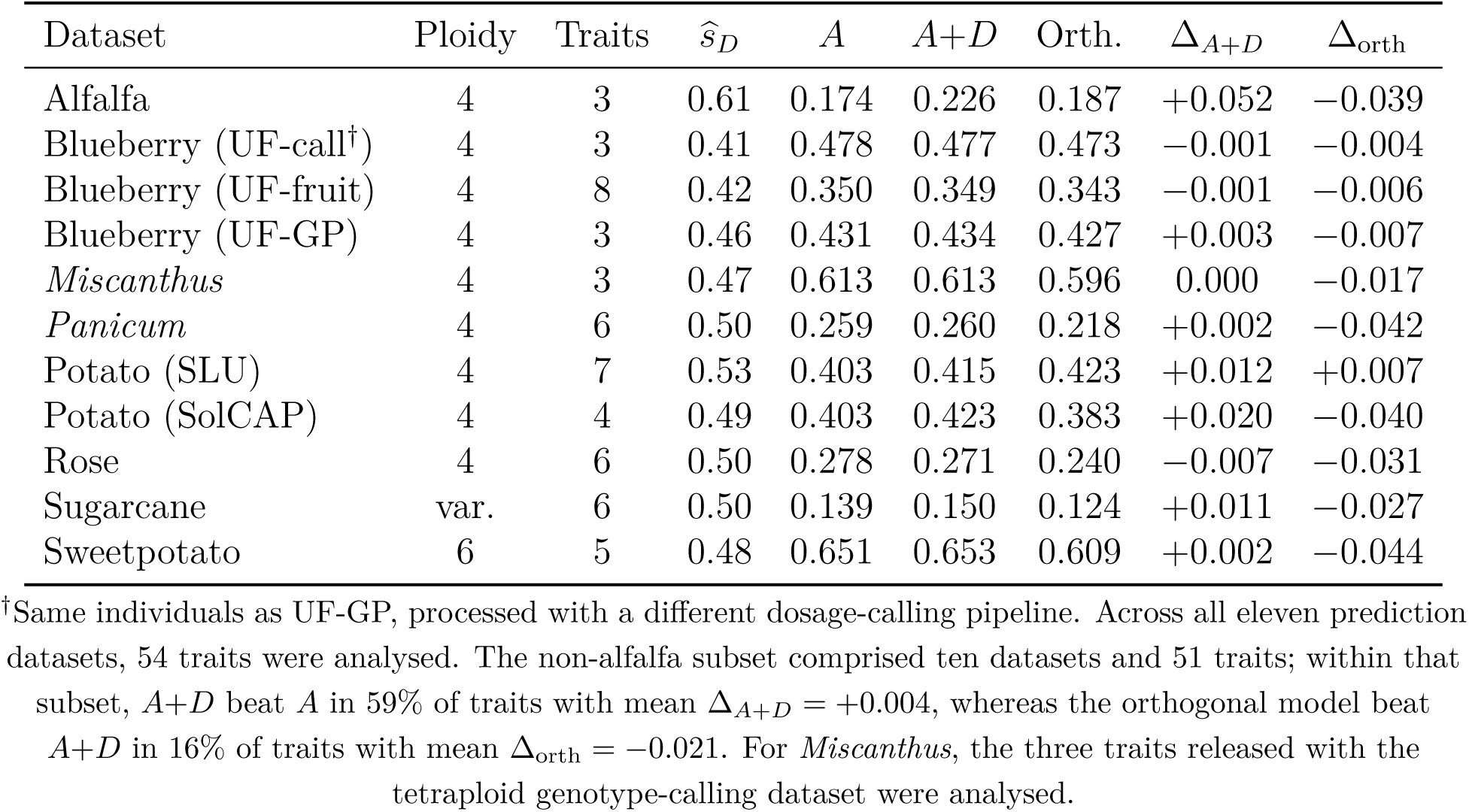
Cross-crop predictive ability of additive, standard *A* + *D* and orthogonal dosage-degree models. For each dataset, the table gives ploidy, number of traits, the standard *D*-kernel share from the full-data diagnostic fits, *s_D_* = *V*_D_*/*(*V*_A_ + *V*_D_), averaged over each dataset’s diagnostic traits (for alfalfa, its single representative yield trait), mean cross-validated predictive ability for additive-only (*A*), standard *A* + *D* and orthogonal *A* + *D*_2_ + + *D_m_* models, and paired prediction gains Δ*_A_*_+_*_D_* = (*A*+*D*) *A* and Δ_orth_ = orth (*A*+*D*). All rows use paired five-fold, five-repeat cross-validation with identical fold partitions across models. Trait-mean values are descriptive.

| Dataset | Ploidy | Traits | *ŝ*<sub>*D*</sub> | *A* | *A*+*D* | Orth. | Δ<sub>*A*+*D*</sub> | Δ<sub>orth</sub> |
|---|---|---|---|---|---|---|---|---|
| Alfalfa | 4 | 3 | 0.61 | 0.174 | 0.226 | 0.187 | +0.052 | −0.039 |
| Blueberry (UF-call<sup>†</sup>) | 4 | 3 | 0.41 | 0.478 | 0.477 | 0.473 | −0.001 | −0.004 |
| Blueberry (UF-fruit) | 4 | 8 | 0.42 | 0.350 | 0.349 | 0.343 | −0.001 | −0.006 |
| Blueberry (UF-GP) | 4 | 3 | 0.46 | 0.431 | 0.434 | 0.427 | +0.003 | −0.007 |
| *Miscanthus* | 4 | 3 | 0.47 | 0.613 | 0.613 | 0.596 | 0.000 | −0.017 |
| *Panicum* | 4 | 6 | 0.50 | 0.259 | 0.260 | 0.218 | +0.002 | −0.042 |
| Potato (SLU) | 4 | 7 | 0.53 | 0.403 | 0.415 | 0.423 | +0.012 | +0.007 |
| Potato (SolCAP) | 4 | 4 | 0.49 | 0.403 | 0.423 | 0.383 | +0.020 | −0.040 |
| Rose | 4 | 6 | 0.50 | 0.278 | 0.271 | 0.240 | −0.007 | −0.031 |
| Sugarcane | var. | 6 | 0.50 | 0.139 | 0.150 | 0.124 | +0.011 | −0.027 |
| Sweetpotato | 6 | 5 | 0.48 | 0.651 | 0.653 | 0.609 | +0.002 | −0.044 |

<sup>†</sup>Same individuals as UF-GP, processed with a different dosage-calling pipeline. Across all eleven prediction datasets, 54 traits were analysed. The non-alfalfa subset comprised ten datasets and 51 traits; within that subset, *A*+*D* beat *A* in 59% of traits with mean Δ<sub>*A*+*D*</sub> = +0.004, whereas the orthogonal model beat *A*+*D* in 16% of traits with mean Δ<sub>orth</sub> = −0.021. For *Miscanthus*, the three traits released with the tetraploid genotype-calling dataset were analysed.

A large standard *D*-kernel share was not sufficient for predictive gain. Although the standard *A* + *D* model assigned a substantial share of fitted variance to the *D* kernel in every prediction dataset, adding this kernel to the additive model usually produced little or no cross-validated improvement (Figure 5A). Alfalfa was the clearest exception, with a consistent gain for biomass-yield prediction. The two potato datasets and sugarcane showed smaller positive gains, whereas blueberry, *Panicum*, rose, *Miscanthus* and sweet-potato showed negligible gains on average. Because some datasets contributed many more traits than others, we emphasize paired contrasts at both the dataset and trait levels rather than relying on a single pooled mean.

These prediction results agree with earlier reports. Amadeu et al. (2020) found similar predictive ability for additive and additive-plus-dominance models in autotetraploid blue-berry and potato, and Song et al. (2024) found no gain from adding dominance for most traits in blueberry, guinea grass and sweetpotato. In the three datasets from the same public sources as the latter study (blueberry UF-GP, *Panicum* and sweetpotato), our paired gains from adding the standard *D* kernel were +0.003, +0.002 and +0.002 (Table 1). The two analyses differ in dominance kernel, estimator and phenotype processing, so this is agreement in the conclusion and not a replication of their estimates.

The orthogonal degree-kernel expansion did not generally improve prediction relative to the standard *A* +*D* model (Figure 5B). Replacing the standard *D* kernel with the orthogonal *A* + *D*_2_ + · · · + *D_m_* model usually reduced or left unchanged predictive ability. This result should not be interpreted as evidence that higher-order dosage effects are biologically absent. Rather, it shows that, in these finite empirical datasets, the additional degree kernels did not provide enough independent predictive signal to offset the extra estimation burden. This pattern is consistent with the kernel-geometry results: the weak empirical separability means that adding them as separate prediction kernels does not necessarily improve out-of-sample accuracy.

Thus, dominance-related kernels should be judged separately for prediction and interpretation. A standard *D* kernel may be useful when it improves validated prediction in the target population, but such a gain demonstrates predictive utility rather than biological identifiability of digenic dominance. Conversely, a large fitted *D*-kernel share does not imply that adding a dominance kernel will materially improve prediction. We therefore recommend reporting cross-validated prediction gains and fitted variance fractions as separate results: retain dominance-related kernels when paired cross-validation supports them, but interpret their fitted variance components biologically only when the identifiability diagnostics also support that interpretation.

### 3.5 Panel-conditioned workflow classifications

We applied the panel-conditioned diagnostic workflow to assign each fitted dominance-labelled component to a reporting category: structure-sensitive, *D*_2_-supported, prediction-useful, or model-basis/unresolved (Figure 6). The workflow was used as an interpretation rule, not as an additional model-fitting procedure: biological *D*_2_ support required excess over the permutation null, recoverability of a planted *D*_2_ signal in the same genotype matrix, localization of the real-trait signal toward the orthogonal *D*_2_ direction and limited posterior uncertainty, whereas prediction gain was evaluated separately.

**Figure 6:**
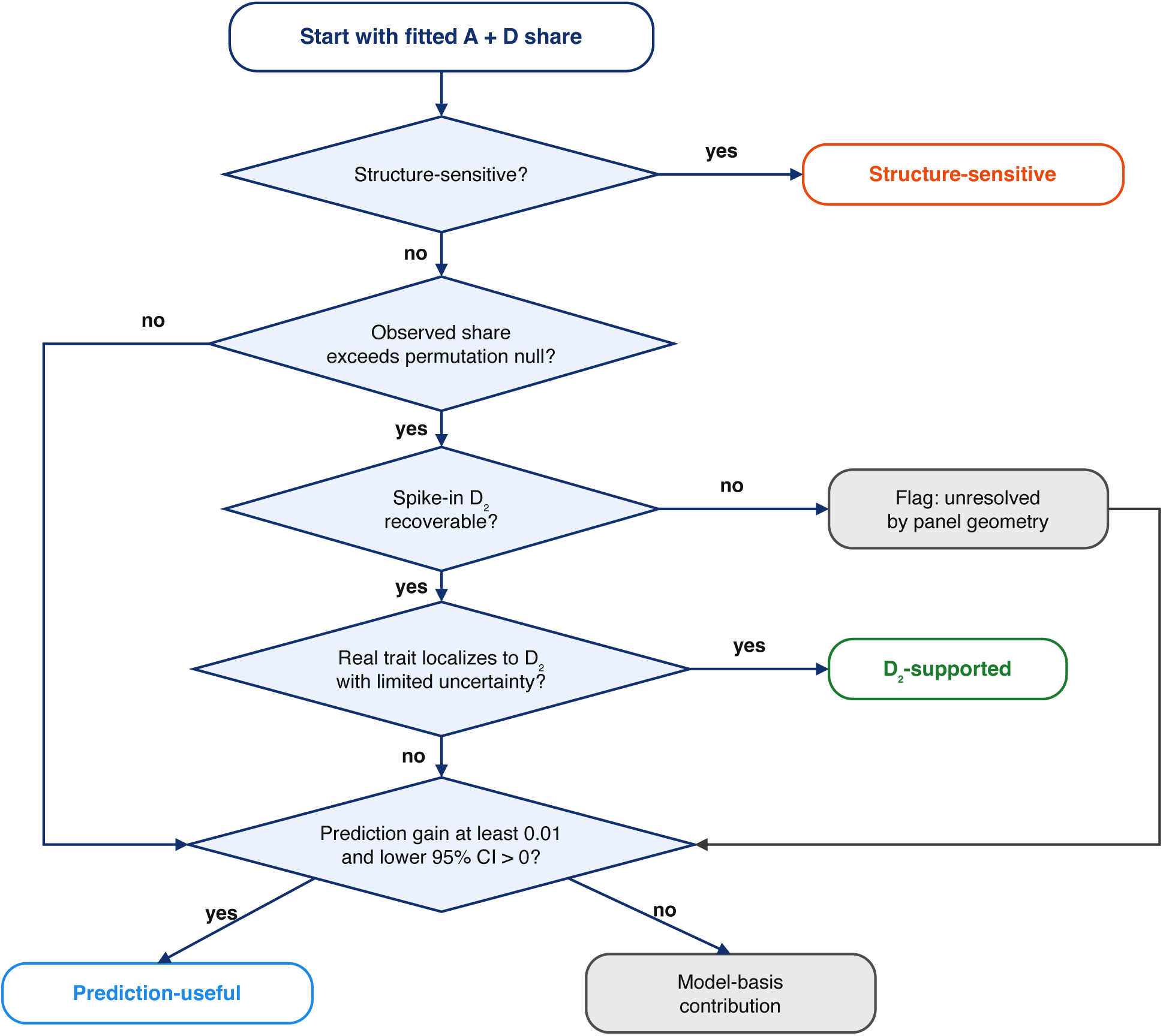
Panel-conditioned diagnostic workflow for dominance-labelled variance. Each panel is analysed with its own genotype matrix, kernels and reference controls rather than against a universal simulation ruler. Step 1 identifies structure-sensitive fitted *D* shares using PC and family/group control calibrated by a structure-spike ruler. Step 2 tests whether the signal is *D*_2_-supported, requiring excess over the permutation null, recoverability of a planted *D*_2_ spike-in, real-trait localization toward *D*_2_ and limited posterior uncertainty. Step 3 separates prediction utility from biological interpretation using paired cross-validation. Prediction gain is the absolute change in held-out Pearson correlation, Δ*_A_*_+_*_D_* = *r*(*A* + *D*) *r*(*A*); prediction-useful requires Δ*_A_*_+_*_D_* 0.01 with the lower 95% confidence bound above zero, the bound being computed over trait-by-repeat contrasts (Supplemental Table S6). If the planted *D*_2_ spike-in is not recoverable, the result is flagged as unresolved by panel geometry but still proceeds to this prediction-gain test. Biological digenic dominance is claimed only when all Step 2 criteria pass; prediction gain alone supports retaining the kernel for prediction but never licenses a biological *D*_2_ claim.

No empirical analysis was classified as *structure-sensitive* or *D*_2_*-supported*. Structure-control changes were small, indicating that the fitted standard *D* components were not simply removed by genomic principal-component adjustment or by available family, cohort or group adjustment. However, this robustness did not make the fitted components biologically interpretable. Observed standard *D*-kernel shares exceeded the panel-specific no-signal reference by more than the margin in only two panels (alfalfa and potato SLU), and in those two the localization criteria were not met; in all panels the workflow’s Bayesian orthogonal fits did not localize non-additive variance to *D*_2_, and posterior intervals for non-additive degree fractions were broad or overlapping. Full decision variables for each dataset are reported in Supplemental Table S6.

The absence of empirical *D*_2_ support was generally not explained solely by failure of the genotype matrices to recover planted signal. In several panels, the *D*_2_ spike-in control was recoverable, but the corresponding real traits did not localize toward *D*_2_. Thus, the limiting step was usually the real-trait evidence for *D*_2_ localization, not only the panel’s theoretical capacity to detect a planted *D*_2_ component.

Prediction gain gave a separate classification. Panels with a material paired cross-validation gain from adding the standard *D* kernel were labelled *prediction-useful*; this supported retaining the dominance-labelled kernel for prediction but not interpreting its fitted variance as biological digenic dominance. Panels without material prediction gain were labelled *model-basis*. Under either label, a panel carried an additional unresolved-by-geometry flag when the planted *D*_2_ spike-in was not recoverable.

The *model-basis* label is not evidence that digenic dominance is absent. It records that the criteria for support were not met, which can also result from weak signal, limited power, model misspecification or an architecture that is not a single dosage degree. Panels in which the planted *D*_2_ signal was not recovered carry the unresolved-by-geometry flag for this reason. A panel without the flag has shown recoverability only for the planted setting, a digenic fraction of 0.5 at *H*^2^ = 0.6, so a smaller digenic contribution could remain undetected in it.

The workflow did not reject biological interpretation by construction. We passed a separable *D*_2_ positive-control simulation (1,500 individuals, 2,500 independent loci, purely digenic effects, *H*^2^ = 0.6) through every stage of the workflow as a twelfth panel, with the same settings and thresholds as the empirical panels (Supplemental Table S6). It met every criterion. The standard share exceeded its permutation null by 0.27. The planted spike-in was recovered in all replicates. The fitted *D*_2_ fraction (0.74) was closer to the spike-in reference than to the additive-only reference, and its excess over the higher degrees was above all 30 additive-only replicates. The credible-interval width was 0.14, and structure sensitivity was zero. The control was therefore classified *D*_2_*-supported* under the full rule, and it kept that label when the thresholds were varied by 20%. This shows that the criteria can license a biological interpretation when both panel geometry and trait signal support it. The control is an idealized case: its additive–dominance kernel cosine was 0.77, lower than in any empirical panel (0.90 to 0.99), and all of its genetic variance was digenic. It does not show that the workflow has adequate power for the weaker and mixed architectures expected in real traits, and no empirical panel provides a real-data example of a *D*_2_*-supported* result.

## 4 Discussion

### 4.1 Fitted dominance shares should be interpreted according to the claim

This study shows that fitted dominance-related variance in dosage-based polyploid genomic prediction should be interpreted according to the claim being made. A standard *D* kernel can be useful for prediction, and its fitted share can be substantial in fitted models. However, these properties do not by themselves make *V*_D_*/*(*V*_A_ + *V*_D_) a calibrated estimate of biological digenic dominance. In known-truth simulations, the standard *D* share was positive when true *D*_2_ variance was zero and responded weakly to increasing planted *D*_2_ signal. In empirical autotetraploid panels, observed shares did not exceed phenotype-permutation no-signal references by more than the margin in seven of nine datasets, additive and dominance-related kernels were highly collinear, and default-prior orthogonal decompositions did not localize fitted non-additive variance to *D*_2_ (Figures 1, 2 and 4). The safest default interpretation is therefore that the standard *D* share is a dominance-labelled model-basis contribution unless panel-conditioned diagnostics support a stronger biological claim.

Earlier studies had already reported similar predictive ability with and without dominance in blueberry and potato (Amadeu et al., 2020), and no improvement for most traits in blueberry, guinea grass and sweetpotato, with blueberry yield as an exception (Song et al., 2024). What a prediction comparison cannot show is how to read the fitted share. Song et al. interpreted the absence of gain for most traits as dominance variance not being captured (Song et al., 2024), yet the standard *A* + *D* model fitted here assigned between 0.46 and 0.50 of the fitted genetic variance to the *D* kernel in the three datasets from the same sources (Table 1). The two statements are compatible once the share is calibrated: in the simulations a share of this size arose with no digenic dominance, and in these datasets it was close to or below the share returned by the same model for permuted phenotypes. The simulations and panel-conditioned controls therefore add the reference values against which a reported share can be judged, which the earlier prediction studies did not provide.

The fitted *D*-kernel variance was also not partitioned into dominance and family co-variance. Under random mating, without inbreeding or epistasis, the genetic covariance between full sibs has the classical expectation 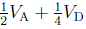 for a diploid, and the corresponding autotetraploid expectation also contains trigenic and quadrigenic terms (Gallais, 2003; Kempthorne, 1957). With recorded families, such expectations allow dominance to be separated from other sources of family resemblance. We did not attempt this partition. All analyses here used marker-based kernels, and the family, cohort and group controls in the workflow test sensitivity to structure without replacing a pedigree-based partition. Some source populations do contain full-sib families, for example the blueberry breeding population and the sugarcane progeny, so a pedigree-based check is possible there and is left for future work.

### 4.2 Finite-panel kernel separability limits calibration and localization

The orthogonal contrast framework clarifies why this happens. Additive, digenic and higher-order intralocus contrasts are well defined at the single-locus reference-population level, and positive-control simulations show that degree-specific signal can be recovered when the causal architecture and genotype design provide sufficient separation. The obstacle in realistic panels is empirical rather than definitional. Linkage disequilibrium, relatedness, allele-frequency imbalance, dosage uncertainty and finite sample size can compress genome-wide kernels into a low-dimensional span. Consequently, population-level orthogonality of dosage contrasts does not guarantee sample-level separability of variance components. Sensitivity analyses sharpen this point: in finite panels, empirical separability of the fitted kernel basis governs calibration. Poor separation between additive and digenic directions produces a positive no-*D*_2_ baseline and attenuated response to true *D*_2_ signal, whereas better-separated kernel bases yield more accurate *D*_2_ estimation.

### 4.3 Prediction gain and structure robustness do not establish biological dominance

The empirical results should therefore be read as a limitation on interpretation, not as evidence that non-additive biology is absent. Alfalfa biomass yield illustrates the distinction: the standard *D* kernel improved cross-validated prediction, but the fitted variance did not satisfy the diagnostic criteria for biological *D*_2_ support. Most other panels showed similar fitted *D* shares but little predictive gain (Figure 5). Thus, prediction and biological interpretation should be reported as separate outcomes. A dominance-related kernel should be retained when paired validation shows useful prediction gain in the target breeding population, but that gain should not be used as evidence for digenic dominance.

Structure-control analyses were consistent with this interpretation. In the empirical work-flow, no real analysis was classified as structure-sensitive: fitted standard *D*-kernel shares were not substantially reduced by genomic principal-component adjustment or by available family, cohort or group adjustment. This weakens a simple structure-only explanation, but it does not make the fitted component biologically interpretable. The remaining diagnostics still pointed to weak identification: observed shares did not exceed dataset-specific no-signal references in most datasets, the Bayesian orthogonal fits did not localize non-additive variance to *D*_2_, posterior intervals for non-additive degrees were broad or overlapping, and additive–dominance kernels remained poorly separable. Persistence after structure adjustment should therefore be reported as robustness of a fitted kernel contribution, not as evidence for biological digenic dominance.

### 4.4 Diagnostic reporting and next steps

The practical implication is diagnostic rather than dismissive (Figure 6). We recommend reporting standard *D*-kernel shares together with other diagnostic metrics. Biological digenic dominance should be claimed only when these diagnostics align: the observed signal exceeds the no-signal null, the relevant kernels are empirically separable, planted *D*_2_ signal is recoverable in the same genotype matrix, the real trait localizes to *D*_2_ with limited un-certainty, and the conclusion is robust to structure control. When these conditions fail, the more accurate wording is that the dominance-labelled kernel explains a share of fitted model variance.

The workflow and its labels rest on one estimator, Bayesian kernel regression in BGLR (Pérez & de los Campos, 2014) with its default priors, and the estimator comparison shows what this choice does (Supplemental Table S5). When kernels are highly collinear the data constrain the split of genetic variance among them only weakly: REML intervals for the standard share were wide, and the Bayesian fit resolved the weak constraint toward shares near one-half and near-equal *D*_2_, *D*_3_ and *D*_4_ fractions. Those Bayesian fractions should not be read as estimates of the degree composition of non-additive variance. The two approaches overlapped only partly in where they indicated a dominance-labelled signal. The REML intervals excluded a zero share in alfalfa and the two potato panels and for two blueberry UF-GP traits. Of these, alfalfa and potato SLU exceeded the permutation null, and alfalfa and both potato panels were labelled *prediction-useful*; sugarcane was labelled *prediction-useful* without REML support, and blueberry UF-GP was labelled *model-basis*. The estimators also disagreed on composition, with REML assigning the non-additive variance to *D*_2_. Because every reference control in the workflow passes through the same Bayesian estimator, the labels are internally consistent, but a workflow built on REML could label alfalfa and potato differently. This was not evaluated: REML localization was not calibrated against known truth, and the REML degree fractions have no intervals. Known-truth calibration of likelihood-based localization is the first next step.

Several limitations define next steps. The reference distributions are small: 30 permutations per trait and 30 replicates per control cell, so the additive-only percentile used for localization resolves steps of 1*/*30, and the decision thresholds are conventions rather than calibrated error rates (their sensitivity is reported in Supplemental Section C.5). Double reduction was not modelled in the AlphaSimR simulations, the positive control or the contrast basis. It moves genotype frequencies away from the binomial reference, so the contrasts are then not orthogonal even at the population level. An approximation in a separate sensitivity grid, which replaced heterozygous dosages by homozygous ones with probability 0.1, left the additive-truth standard share large (0.39 against 0.42) and lowered the effective rank of the degree kernels (Supplemental Methods); a meiosis-based model of double reduction was not tested. The empirical analyses used public datasets that differ in marker density, dosage-calling quality, allele-frequency spectra, phenotype precision and population design, rather than a balanced survey of all autopolyploid breeding systems. The positive controls show that the workflow can identify interpretable *D*_2_ signal, but they do not exhaust all possible non-additive architectures. Future work should prioritize designed mating populations, phased haplotypes where available, explicit double-reduction modelling, empirical-frequency orthogonalization, per-trait permutation and spike-in calibration, likelihood profiling and multi-environment models that separate dominance, heterozygosity, epistasis and genotype-by-environment interaction.

Overall, dominance kernels remain useful tools for polyploid genomic prediction, especially in clonal, heterotic or family-structured breeding contexts. What should change is the language used for their variance components: unless explicit diagnostics support biological identifiability, a fitted dominance share should be reported as a dominance-labelled kernel contribution rather than as an estimate of biological digenic dominance.

## Supplemental Material

The supplemental material contains Supplemental Methods, supplemental simulation results and empirical-panel diagnostics, Supplemental Tables S1–S7 and Supplemental Figure S1.

## Data Availability Statement

All empirical panels analysed here are public: alfalfa Roza2019 (Medina et al., 2025); potato SolCAP (Rosyara et al., 2016); potato SLU (Selga et al., 2022); blueberry (de Bem Oliveira et al., 2020; Ferrão et al., 2018; Njuguna et al., 2023); *Panicum maximum* (Lara et al., 2019); rose (Wamhoff et al., 2024); *Miscanthus* (Njuguna et al., 2023); and sweetpotato and sugarcane (Batista et al., 2022). The code for the orthogonal dosage-contrast framework and the diagnostic workflow, polyDx, is open source at https://github.com/hirotong/ polyploid-dominance-identifiability, together with the scripts and the summary tables that produce every figure and table in this article.

## Conflict of Interest Statement

The authors declare no conflicts of interest.

## Author Contributions

Jinguang Tong: Conceptualization; Methodology; Software; Formal analysis; Investigation; Visualization; Writing – original draft. Qingchuan Yang: Conceptualization; Supervision; Writing – review & editing. Mingna Li: Conceptualization; Supervision; Writing – review & editing.

## Acknowledgments

This work was supported by the Central Public-interest Scientific Institution Basal Research Fund (No. 2026-YWF-ZYSQ-04S) and the Hohhot Science and Technology Plan Project (2024-Unveiled and Leading-agriculture-2-2).

## Abbreviations

*D*_2_: digenic dominance
LD: linkage disequilibrium
NOIA: natural and orthogonal interactions
PC: principal component
REML: restricted maximum likelihood
RMSE: root mean squared error
SNP: single-nucleotide polymorphism

## Notes

### Competing Interest Statement

The authors have declared no competing interest.

## References

1. Álvarez-Castro, J. M., & Carlborg, Ö. (2007). A unified model for functional and statistical epistasis and its application in quantitative trait loci analysis. Genetics, 176 (2), 1151– 1167. 10.1534/genetics.106.067348

2. Amadeu, R. R., Ferrão, L. F. V., de Bem Oliveira, I., Benevenuto, J., Endelman, J. B., & Muñoz, P. R. (2020). Impact of dominance effects on autotetraploid genomic prediction. Crop Science, 60 (2), 656–665. 10.1002/csc2.20075

3. Amadeu, R. R., Garcia, A. A. F., Munoz, P. R., & Ferrão, L. F. V. (2023). AGHmatrix: genetic relationship matrices in R. Bioinformatics, 39 (7), btad445. 10.1093/bioinformatics/btad445

4. Batista, L. G., Mello, V. H., Souza, A. P., & Margarido, G. R. A. (2022). Genomic prediction with allele dosage information in highly polyploid species. Theoretical and Applied Genetics, 135 (2), 723–739. 10.1007/s00122-021-03994-w

5. Chen, J., Zhang, F., Wang, L., Leach, L., & Luo, Z. (2018). Orthogonal contrast based models for quantitative genetic analysis in autotetraploid species. New Phytologist, 220 (1), 332–346. 10.1111/nph.15284

6. Cockerham, C. C. (1954). An extension of the concept of partitioning hereditary variance for analysis of covariances among relatives when epistasis is present. Genetics, 39 (6), 859–882. 10.1093/genetics/39.6.859

7. de Bem Oliveira, I., Amadeu, R. R., Ferrão, L. F. V., & Muñoz, P. R. (2020). Optimizing whole-genomic prediction for autotetraploid blueberry breeding. Heredity, 125 (6), 437–448. 10.1038/s41437-020-00357-x

8. Endelman, J. B., Schmitz Carley, C. A., Bethke, P. C., Coombs, J. J., Clough, M. E., da Silva, W. L., De Jong, W. S., Douches, D. S., Frederick, C. M., Haynes, K. G., Holm, D. G., Miller, J. C., Muñoz, P. R., Navarro, F. M., Novy, R. G., Palta, J. P., Porter, G. A., Rak, K. T., Sathuvalli, V. R., … Yencho, G. C. (2018). Genetic variance partitioning and genome-wide prediction with allele dosage information in autotetraploid potato. Genetics, 209 (1), 77–87. 10.1534/genetics.118.300685

9. Ferrão, L. F. V., Benevenuto, J., de Bem Oliveira, I., Cellon, C., Olmstead, J., Kirst, M., Resende, M. F. R., Jr., & Muñoz, P. (2018). Insights into the genetic basis of blueberry fruit-related traits using diploid and polyploid models in a GWAS context. Frontiers in Ecology and Evolution, 6, 107. 10.3389/fevo.2018.00107

10. Gallais, A. (2003). Quantitative genetics and breeding methods in autopolyploid plants. Paris: INRA Editions.

11. Gaynor, R. C., Gorjanc, G., & Hickey, J. M. (2021). AlphaSimR: an R package for breeding program simulations. G3: Genes, Genomes, Genetics, 11 (2), jkaa017. 10.1093/g3journal/jkaa017

12. Kempthorne, O. (1957). An introduction to genetic statistics. New York: John Wiley & Sons.

13. Lara, L. A. d. C., Santos, M. F., Jank, L., Chiari, L., Vilela, M. d. M., Amadeu, R. R., dos Santos, J. P. R., Pereira, G. d. S., Zeng, Z.-B., & Garcia, A. A. F. (2019). Genomic selection with allele dosage in Panicum maximum Jacq. G3: Genes, Genomes, Genetics, 9 (8), 2463–2475. 10.1534/g3.118.200986

14. Lopes, M. S., Bastiaansen, J. W. M., Janss, L., Knol, E. F., & Bovenhuis, H. (2015). Estimation of additive, dominance, and imprinting genetic variance using genomic data. G3: Genes, Genomes, Genetics, 5 (12), 2629–2637. 10.1534/g3.115.019513

15. Medina, C. A., Hansen, J., Crawford, J., Viands, D., Sapkota, M., Xu, Z., Peel, M. D., & Yu, L.-X. (2025). Genome-wide association and genomic prediction of alfalfa (Medicago sativa L.) biomass yield under drought stress. International Journal of Molecular Sciences, 26 (2), 608. 10.3390/ijms26020608

16. Muñoz, P. R., Resende, M. F. R., Jr., Gezan, S. A., Resende, M. D. V., de los Campos, G., Kirst, M., Huber, D., & Peter, G. F. (2014). Unraveling additive from non-additive effects using genomic relationship matrices. Genetics, 198 (4), 1759–1768. 10.1534/genetics.114.171322

17. Njuguna, J. N., Clark, L. V., Lipka, A. E., Anzoua, K. G., Bagmet, L., Chebukin, P., Dwiyanti, M. S., Dzyubenko, E., Dzyubenko, N., Ghimire, B. K., Jin, X., Johnson, D. A., Kjeldsen, J. B., Nagano, H., de Bem Oliveira, I., Peng, J., Petersen, K. K., Sabitov, A., Seong, E. S., … Sacks, E. J. (2023). Impact of genotype-calling methodologies on genome-wide association and genomic prediction in polyploids. The Plant Genome, 16 (4), e20401. 10.1002/tpg2.20401

18. Pérez, P., & de los Campos, G. (2014). Genome-wide regression and prediction with the BGLR statistical package. Genetics, 198 (2), 483–495. 10.1534/genetics.114.164442

19. Rosyara, U. R., De Jong, W. S., Douches, D. S., & Endelman, J. B. (2016). Software for genome-wide association studies in autopolyploids and its application to potato. The Plant Genome, 9 (2), plantgenome2015.08.0073. 10.3835/plantgenome2015.08.0073

20. Selga, C., Reslow, F., Pérez-Rodríguez, P., & Ortiz, R. (2022). The power of genomic estimated breeding values for selection when using a finite population size in genetic improvement of tetraploid potato. G3: Genes, Genomes, Genetics, 12 (1), jkab362. (Data: figshare 10.25387/g3.14376938) 10.1093/g3journal/jkab362

21. Slater, A. T., Cogan, N. O. I., Forster, J. W., Hayes, B. J., & Daetwyler, H. D. (2016). Improving genetic gain with genomic selection in autotetraploid potato. The Plant Genome, 9 (3), plantgenome2016.02.0021. 10.3835/plantgenome2016.02.0021

22. Song, H., Zhang, Q., & Hu, H. (2024). polyGBLUP: a modified genomic best linear unbiased prediction improved the genomic prediction efficiency for autopolyploid species. Briefings in Bioinformatics, 25 (2), bbae106. 10.1093/bib/bbae106

23. Vitezica, Z. G., Legarra, A., Toro, M. A., & Varona, L. (2017). Orthogonal estimates of variances for additive, dominance, and epistatic effects in populations. Genetics, 206 (3), 1297–1307. 10.1534/genetics.116.199406

24. Vitezica, Z. G., Varona, L., & Legarra, A. (2013). On the additive and dominant variance and covariance of individuals within the genomic selection scope. Genetics, 195 (4), 1223–1230. 10.1534/genetics.113.155176

25. Wamhoff, D., Marxen, A., Acharya, B., Grzelak, M., Debener, T., & Winkelmann, T. (2024). Genotypic and phenotypic raw data for GWAS study Wamhoff et al. 2024. (Research Data Repository, Leibniz Universität Hannover (LUIS)) 10.25835/xvdntbqg

26. Yadav, S., Wei, X., Joyce, P., Atkin, F., Deomano, E., Sun, Y., Nguyen, L. T., Ross, E. M., Cavallaro, T., Aitken, K. S., Hayes, B. J., & Voss-Fels, K. P. (2021). Improved genomic prediction of clonal performance in sugarcane by exploiting non-additive genetic effects. Theoretical and Applied Genetics, 134 (7), 2235–2252. 10.1007/s00122-021-03822-1

