## Supplemental Material for "Limited biological identifiability of dominance-labelled variance components in polyploid genomic prediction"

### A Supplemental Methods

#### A.1 Empirical-panel sources and preprocessing

The main text gives a compact overview of the empirical genotype–phenotype panels. Here we provide the panel-specific processing details needed to interpret Table S1 and the downstream diagnostic analyses.

We gathered 11 empirical panels across eight crops. The autotetraploid analyses used alfalfa Roza2019 (Medina et al., 2025), potato SolCAP (Rosyara et al., 2016), potato SLU (Selga et al., 2022), blueberry UF-GP (de Bem Oliveira et al., 2020), blueberry UF-fruit (Ferrão et al., 2018), *Panicum maximum* (Lara et al., 2019), rose (Wamhoff et al., 2024) and *Miscanthus* (Njuguna et al., 2023). Higher-ploidy case studies used published autohexaploid sweetpotato and variable-ploidy sugarcane panels (Batista et al., 2022). The blueberry UF-call dataset (Njuguna et al., 2023) was not treated as an independent biological population because it contains the same individuals as the blueberry UF-GP panel but uses an alternative dosage-calling pipeline; we therefore used it as a dosage-calling robustness check.

All panels were analysed on environment-adjusted phenotypes. Published BLUEs, BLUPs, least-squares means or trait summaries were used where the public release provided them, so that the genomic models were fitted to the same phenotype scale used in the original studies whenever possible. For the two panels distributed as per-trial records, potato SLU with multi-location plot data and sweetpotato with multi-year records, we fitted a fixed environment effect, using location or year as appropriate, and used the resulting genotype entry means as BLUEs. This avoided confounding genotype with environment in the unbalanced trial designs. For multi-trait panels, analyses were conducted trait by trait using the individuals with non-missing phenotype records for that trait. Dataset-level summaries were then computed by averaging over analysed traits unless otherwise stated. For alfalfa, the full-length variance-component summaries and the localization diagnostics used the overall biomass-yield BLUP as the representative trait, whereas the permutation null, the estimator comparison and the prediction summaries used the three closely related biomass-yield traits. For *Miscanthus*, analyses used the three least-squares-mean traits released with the tetraploid genotype-calling dataset.

Dosage matrices were used as published except for blueberry UF-GP, for which we called dosages from the published read counts with `updog` (Gerard et al., 2018) rather than using the genotype calls of the source study. For sugarcane, all markers were retained at their native per-marker ploidy (in $\{6, 8, 10, 12, 14\}$) and only the common quality-control filters were applied. Because the sugarcane data represent variable-ploidy allele counts, the diagnostic degree-kernel analysis used contrasts computed at each marker's own ploidy but truncated the orthogonal decomposition to a fixed maximum degree of six.

For each panel, individuals were retained after matching genotype records to the available phenotypes. A common marker quality-control procedure was then applied across panels. Markers were retained if they were polymorphic, had minor-allele frequency at least 0.05 and had call rate greater than 0.8. Remaining missing dosages were imputed by the marker mean before kernel construction. Dosage covariates were centred and scaled within each empirical panel after genotype–phenotype alignment, so all kernels were constructed from the realized marker matrix available to the corresponding fitted model. Sample sizes, marker counts after filtering, analysed traits, ploidy levels, phenotype summaries and genotype sources are reported in Table S1.

**Table S1: Empirical polyploid datasets analysed.** Shown are crop, public dataset identifier (with its literature reference), ploidy, sample size after genotype–phenotype alignment ($n$), marker count after the common quality-control filters (polymorphic loci with minor-allele frequency $\geq 0.05$ and call rate $> 0.8$), number of analysed traits, a brief phenotype summary and the genotype source. All panels use environment-adjusted phenotypes: published BLUEs, BLUPs or LS-means where released, and genotype entry means (BLUEs) from a fitted environment effect for the potato SLU (multi-location) and sweetpotato (multi-year) panels distributed as per-trial records. The blueberry UF-call panel uses the same individuals as UF-GP with an alternative dosage-calling pipeline and was treated as a robustness check rather than as an independent biological population.

| Crop | Dataset | Ploidy | $n$ | Markers | Traits | Phenotype summary | Genotype source |
|---|---|---|---|---|---|---|---|
| Alfalfa | Roza2019 (Medina et al., 2025) | 4 | 424 | 50,366 | 3 | Biomass-yield summaries | Published GBS dosage (NGSEP, ploidy 4) |
| Blueberry | UF-call† (Njuguna et al., 2023) | 4 | 1,804 | 68,768 | 3 | Fruit weight, firmness, scar diameter | Published `updog` discrete calls |
| Blueberry | UF-fruit (Ferrão et al., 2018) | 4 | 1,559 | 80,591 | 8 | Eight fruit-quality traits | Published discrete dosage |
| Blueberry | UF-GP (de Bem Oliveira et al., 2020) | 4 | 1,804 | 32,319 | 3 | Fruit weight, firmness, yield (LS-means) | Our `updog` calls from published read counts |
| *Miscanthus* | Biomass (Njuguna et al., 2023) | 4 | 197 | 71,431 | 3 | Culm length, biomass yield, heading date | Published `updog` discrete calls |
| *Panicum* | Forage (Lara et al., 2019) | 4 | 530 | 14,456 | 6 | Forage-quality traits | Published dosage |
| Potato | SLU (Selga et al., 2022) | 4 | 669 | 10,545 | 7 | Tuber and quality traits | Published dosage |
| Potato | SolCAP (Rosyara et al., 2016) | 4 | 187 | 3,179 | 4 | Yield, shape, sucrose, maturity | SNP-array ACGT calls |
| Rose | Rooting (Wamhoff et al., 2024) | 4 | 106 | 27,701 | 6 | In-vitro rooting traits | AXIOM SNP-chip dosage |
| Sugarcane | GS (Batista et al., 2022) | 6–14 | 170 | 4,803 | 6 | Brix, Pol, fiber, morphology | Variable-ploidy allele counts |
| Sweetpotato | GS (Batista et al., 2022) | 6 | 282 | 53,802 | 5 | Color traits $a, b, C, h, L$ | Published hexaploid dosage |

†Same individuals as UF-GP, processed with a different dosage-calling pipeline. The alfalfa panel has one genotyped plant from each of 424 half-sib families. The *Panicum* panel is 530 sexual offspring of 19 half-sib families. For *Miscanthus*, the source included 226 tetraploid accessions; our genotype–phenotype alignment retained 197. The sugarcane release contains 168 progeny and two parents; the source study describes 179 progeny.

#### A.2 Simulation implementation

Primary known-truth autotetraploid simulations were generated with AlphaSimR (Gaynor et al., 2021) from a shared founder population so that population-structure comparisons differed in relatedness pattern rather than in marker set or causal architecture. A single coalescent founder pool with 450 founders and 2,400 SNP markers was generated and used to derive three $n = 400$ panels: an unrelated panel sampled from founders, a paternal half-sib panel with 50 families of eight individuals and a full-sib panel with 50 families of eight individuals from random crosses among 40 founders. The same marker and causal-locus definitions were used across the three structures. Meiosis used the AlphaSimR default of bivalent pairing (quadrivalent probability zero), so double reduction did not occur in these populations.

For each population structure, traits were generated directly from the orthogonal dosage-contrast covariates. Pure single-degree architectures placed causal effects only on one degree: additive, digenic, trigenic or quadrigenic. A directional heterozygosity scenario was also included, in which genetic values followed a genome-wide heterozygosity gradient rather than a single orthogonal degree. This scenario was intended as a non-localizing control, because it need not map uniquely onto $D_2$, $D_3$ or $D_4$. Genetic values were standardized within panel, and residual noise was added to target broad-sense heritability $H^2 = 0.6$. The primary pure-architecture simulations used 50 replicates per architecture and population structure.

Calibration simulations used mixed additive-plus-digenic truth to test whether fitted dominance-labelled fractions tracked the known amount of biological $D_2$ signal. In each replicate, standardized additive and digenic genetic values, $G^*_A$ and $G^*_{D_2}$, were generated from the degree-one and degree-two contrast covariates. The genetic value was formed as

$$G = \sqrt{1-s}\, G^*_A + \sqrt{s}\, G^*_{D_2},$$

where

$$s \in \{0, 0.05, 0.10, 0.25, 0.50, 0.75\}.$$

Residual noise was again added to target $H^2 = 0.6$. Because $G^*_A$ and $G^*_{D_2}$ can be empirically correlated in finite panels, the realized $G^*_A$–$G^*_{D_2}$ correlation was recorded per replicate as a diagnostic; the recorded true digenic fraction is the marginal component-variance ratio, which by construction equals the nominal $s$. Additive-only, standard $A + D$ and orthogonal $A + D_2 + D_3 + D_4$ models were fitted for 100 replicates in each structure–fraction cell, and calibration was summarized by comparing fitted dominance-labelled fractions with the nominal true digenic fraction $s$.

The workflow positive control was a separate simulation, not generated with AlphaSimR. Dosages for 1,500 autotetraploid individuals at 2,500 loci were drawn independently from binomial distributions with allele frequencies uniform on $[0.15, 0.85]$, so loci were in Hardy–Weinberg and linkage equilibrium. Genetic values were the sum of normally distributed effects on the degree-two contrast at 300 randomly chosen loci, standardized, with residual noise added to give $H^2 = 0.6$. The simulated genotypes and phenotype were stored as a panel and passed through every stage of the panel-conditioned workflow, with the same number of permutations, control replicates and thresholds as the empirical panels.

Double reduction was examined only through an approximation in a separate additive-truth sensitivity grid ($n \in \{150, 300, 600, 1{,}200\}$; 1,000 or 5,000 markers; two linkage-disequilibrium levels; three population structures; 50 replicates). After genotypes were simulated, each heterozygous dosage was replaced with probability 0.1 by a homozygous dosage, keeping the expected dosage. This is not a model of meiosis. Averaged over the grid, the approximation changed the standard $D$-kernel share under additive truth from 0.42 to 0.39 (all three structures). The kernel geometry was recorded for the full-sib panels only; there the additive–dominance kernel cosine changed from 0.91 to 0.86 and the effective rank of the four orthogonal kernels from 2.21 to 1.70. The share therefore stayed large, and the degree kernels became less separable.

#### A.3 BGLR fitting and posterior summaries

All variance-component models were fitted with `BGLR` (Pérez & de los Campos, 2014) using one RKHS term per genomic relationship kernel. For the additive-only model, the fitted kernel set contained only $K_A$. For the standard model, the fitted kernel set contained $K_A$ and the standard dominance-labelled kernel. For the orthogonal model, the fitted kernel set contained the degree kernels $K_1, \ldots, K_m$, with $K_1$ corresponding to the additive dosage contrast and higher $K_k$ corresponding to intralocus non-additive dosage degrees.

Primary empirical posterior summaries used 12,000 MCMC iterations, 3,000 burn-in iterations and thinning interval five. For these empirical fits, variance fractions were computed at each retained MCMC iteration before summarization, rather than by forming ratios of posterior means. The known-truth simulation summaries used shorter chains (4,000 iterations with 1,000 burn-in; 2,000 with 500 in the stress grid, to limit computing time at sample sizes up to 19,200) and report fractions of posterior-mean variance components. For an empirical fit and a retained sample $t$,

$$s_D^{(t)} = \frac{\sigma_D^{2(t)}}{\sigma_A^{2(t)} + \sigma_D^{2(t)}}$$

for the standard model, and

$$p_k^{(t)} = \frac{\sigma_k^{2(t)}}{\sum_l \sigma_l^{2(t)}}$$

for the orthogonal model. Posterior means were computed from these per-iteration fractions, and 95% credible intervals were computed from their 2.5% and 97.5% posterior quantiles. This calculation preserves posterior dependence among variance components. Simulation summaries are reported as means and standard deviations across simulation replicates, whereas empirical summaries are reported as posterior means with credible intervals where appropriate.

For high-throughput workflow components requiring many repeated fits, including phenotype permutations and selected structure-control checks, a shorter point-estimation run was used with 3,000 MCMC iterations and 750 burn-in iterations. These runs were used for panel-level diagnostic scores rather than for the main posterior interval summaries.

#### A.4 Estimator and prior sensitivity

To see how far the fitted fractions depend on the estimator, the first three analysed traits of every panel (the traits used for the permutation null) were refitted on the same kernels in two ways. First, the standard and orthogonal models were refitted in `BGLR` with the primary chain settings in seven runs per trait: the default priors (prior proportion of variance explained $R^2 = 0.5$ and prior degrees of freedom $\mathrm{df}_0 = 5$ for each variance) with three random seeds, and four changed settings, $R^2 \in \{0.2, 0.8\}$ and $\mathrm{df}_0 \in \{1, 30\}$, each changed alone. Second, both models were fitted by restricted maximum likelihood (REML) with an intercept as the only fixed effect, using individuals with a recorded phenotype. For the standard model, the restricted likelihood was profiled over the share $s_D$ on a grid from zero to one in steps of 0.02: for each value the combined kernel $(1-s_D)K_A+s_DK_D$ was formed and the likelihood was maximised over the ratio of residual to genetic variance. The REML estimate is the grid maximiser, and the 95% interval contains the grid values whose likelihood-ratio statistic against the maximum is at most 3.84; this cut-off is approximate when the estimate is at zero or one. The same profile was computed for 30 permutations of each phenotype. For the orthogonal model, the restricted likelihood was maximised directly over the log variances of all degree kernels and the residual (L-BFGS-B, two starting points), and the fitted variances were converted to fractions; no interval was computed for these fractions. The REML code was checked on simulated data with known shares (script distributed with the released code).

#### A.5 Panel-conditioned diagnostic quantities

The phenotype-permutation null was used to calibrate the standard dominance-labelled share against each panel's own genotype matrix. In the panel workflow, each selected analysed trait was permuted 30 times, the standard $A + D$ model was refit, and the resulting $\widehat{s}_D$ values were recorded. The observed excess over the null was

$$\Delta_{\text{null}} = \widehat{s}_{D,\text{real}} - \overline{\widehat{s}_{D,\text{perm}}}.$$

Observed and null shares were compared like for like. The permuted traits were the first three analysed traits of each panel. For those same traits, $\widehat{s}_{D,\text{real}}$ was computed with the same point estimator as the permutations, the ratio of posterior-mean variance components from the 3,000-iteration run described above, and $\Delta_{\text{null}}$ was averaged over traits. The standard $D$-kernel shares reported elsewhere (Table S4 and the prediction table of the main text) are posterior means from the full-length fits averaged over all analysed traits (for alfalfa, its representative yield trait); they are descriptive and do not enter $\Delta_{\text{null}}$. These quantities were used as descriptive panel-calibration diagnostics, not as formal trait-level hypothesis tests.

Structure sensitivity was summarized as the largest positive reduction in the fitted standard $D$-kernel share after structure control:

$$S_{\text{struct}} = \max_c \left[\widehat{s}_{D,0} - \widehat{s}_{D,c}\right]_+ ,$$

where $[x]_+ = \max(x, 0)$, $\widehat{s}_{D,0}$ is the uncontrolled standard $D$-kernel share, and $\widehat{s}_{D,c}$ is the share under structure-control setting $c$. To calibrate $S_{\text{struct}}$ against each panel's own genotype matrix, a structure-spike ruler repeatedly re-derived $S_{\text{struct}}$ from phenotypes carrying a planted structure-correlated signal of strength $\lambda$. At $\lambda = 0$ (no planted structure effect), the induced $S_{\text{struct}}$ values form the panel's false-positive baseline, and $q_{0.95}^{\text{struct}}$ was defined as the 95th percentile of that baseline distribution, taken as the maximum over structure-control types.

Spike-in recoverability was used to test whether a panel could recover a known planted $D_2$ signal. For each empirical genotype matrix, phenotypes were simulated with a mixed additive-plus-digenic architecture using a planted digenic fraction of $s = 0.5$. The orthogonal model was then refit, and a spike-in replicate was counted as recovered when $D_2$ was the largest fitted non-additive degree. The recovery score was

$$R_{D_2} = \Pr\left(\widehat{p}_{D_2} > \max_{k \geq 3} \widehat{p}_{D_k}\right).$$

For real-trait orthogonal fits, $D_2$ localization was summarized by

$$P_{D_2} = \widehat{p}_{D_2}, \qquad L_{D_2} = P_{D_2} - \max_{k \geq 3} \widehat{p}_{D_k}.$$

The posterior credible-interval width for $D_2$ was recorded as

$$W_{D_2} = Q_{0.975}(p_{D_2}^{(t)}) - Q_{0.025}(p_{D_2}^{(t)}).$$

The median $P_{D_2}$ from the $s = 0.5$ planted-$D_2$ spike-in was denoted $\widetilde{P}_{D_2,\text{spike}}$, and the median $P_{D_2}$ from the additive-only control was denoted $\widetilde{P}_{D_2,\text{add0}}$. The percentile of the observed $L_{D_2}$ within the additive-only control distribution was denoted $q_{L,\text{add0}}$.

### B Supplemental known-truth simulation results

#### B.1 Calibration metrics by structure

To quantify calibration, fitted dominance fractions were regressed on the known digenic fraction, $\widehat{s} = \alpha + \beta s$, separately within each population structure across six values of $s$. Perfect calibration corresponds to $\alpha = 0$ and $\beta = 1$. The resulting intercepts, slopes, RMSEs and endpoint errors are reported in Table S2.

**Table S2: Per-structure calibration of fitted dominance fractions.** Fitted fractions were regressed on the true digenic fraction as $\widehat{s} = \alpha + \beta s$ across six values of $s$ and 100 replicates per value. Perfect calibration has $\alpha = 0$, $\beta = 1$, RMSE = 0 and zero mean error. The standard fitted fraction is $\widehat{s}_D = \widehat{V}_\text{D}/(\widehat{V}_\text{A} + \widehat{V}_\text{D})$; the orthogonal fitted fraction is $\widehat{s}_{D_2}$. $\Delta_0$ and $\Delta_{0.75}$ denote the mean fitted-minus-true difference, $\widehat{s} - s$, at $s = 0$ and $s = 0.75$, respectively.

| | | Calibration fit | | | Mean error | |
|---|---|---|---|---|---|---|
| Fitted fraction | Structure | $\alpha$ | $\beta$ | RMSE | $\Delta_0$ | $\Delta_{0.75}$ |
| $\widehat{s}_D$ | unrelated | 0.39 | 0.26 | 0.28 | +0.37 | −0.17 |
| | half-sib | 0.39 | 0.27 | 0.28 | +0.38 | −0.16 |
| | full-sib | 0.40 | 0.29 | 0.29 | +0.38 | −0.15 |
| $\widehat{s}_{D_2}$ | unrelated | 0.11 | 0.43 | 0.17 | +0.12 | −0.30 |
| | half-sib | 0.11 | 0.40 | 0.18 | +0.13 | −0.32 |
| | full-sib | 0.11 | 0.42 | 0.17 | +0.13 | −0.30 |

#### B.2 Alternative dominance-kernel implementations

To check that the dominance misattribution was not specific to our manual heterozygosity-style $D$ kernel, we repeated the known-truth architectures with two kernels implemented in AGHmatrix (Amadeu et al., 2023): the Slater full-autopolyploid kernel, which includes additive and non-additive effects (Slater et al., 2016), and the Endelman digenic-dominance kernel (Endelman et al., 2018). Each was rescaled to mean diagonal one and fitted alongside the additive kernel in place of the $D$ kernel. Because the Slater kernel also carries additive effects, its share is not a pure dominance share, and its values are shown for comparison with that caveat. The resulting dominance-labelled shares by true architecture are reported in Table S3.

### C Supplemental empirical panel diagnostics

#### C.1 Full posterior summaries for orthogonal degree fractions

Posterior mean fractions and 95% credible intervals for the orthogonal non-additive degrees are reported for the autotetraploid empirical panels in Table S4, together with the standard $D$-kernel share and the kernel-geometry diagnostics.

**Table S3:** Reported $V_{\mathrm{D}}/(V_{\mathrm{A}}+V_{\mathrm{D}})$ by true architecture for three kernels fitted alongside the additive kernel, on the replicate traits of Figure 1A (50 replicates; full-sib panel shown—the unrelated and half-sib panels give the same pattern). The first column is the $D$ kernel of this study and repeats the full-sib values of that figure. Bold values mark the largest spurious shares: the full-autopolyploid Slater kernel under pure higher-order truths, which contain zero true digenic dominance.

| true architecture | manual digenic | Slater | Endelman |
|---|---|---|---|
| additive (true digenic = 0) | 0.37 | 0.33 | 0.24 |
| digenic (control) | 0.65 | 0.72 | 0.73 |
| trigenic (true digenic = 0) | 0.52 | **0.72** | 0.57 |
| quadrigenic (true digenic = 0) | 0.52 | **0.73** | 0.56 |
| directional heterozygosity | 0.70 | 0.53 | 0.46 |

**Table S4: Cross-crop summary of standard $D$-kernel shares, kernel separability and orthogonal degree decomposition.** Shown are the standard $D$-kernel share, $\widehat{s}_D = V_{\mathrm{D}}/(V_{\mathrm{A}} + V_{\mathrm{D}})$, from the standard $A + D$ model; the Frobenius cosine between the additive and standard $D$ kernels; and the effective rank, $r_{\mathrm{eff}}$, of the orthogonal $A + D_2 + D_3 + D_4$ kernel set. The final three columns report posterior mean fractions with 95% posterior credible intervals for the orthogonal non-additive degrees, $\widehat{p}_{D_j} = V_{D_j}/(V_{\mathrm{A}} + V_{D_2} + V_{D_3} + V_{D_4})$ for $j = 2, 3, 4$; the additive fraction is not shown. Observed values are trait-averaged where multiple traits were analysed (posterior means and credible-interval endpoints are arithmetic means of the trait-level values); for alfalfa they are from its single representative yield trait. The comparison with the phenotype-permutation null, on matched traits, is given in Table S6. High $A$–$D$ cosine and low $r_{\mathrm{eff}}$ indicate weak empirical separability of additive and dominance-related kernel directions. Comparable $D_2$, $D_3$ and $D_4$ fractions, together with broad and mutually overlapping credible intervals, indicate that fitted non-additive variance was not localized to the digenic degree.

| Crop | Dataset | $n$ | $\widehat{s}_D$ | A–D cos. / $r_{\mathrm{eff}}$ | $\widehat{p}_{D_2}$ [95% CrI] | $\widehat{p}_{D_3}$ [95% CrI] | $\widehat{p}_{D_4}$ [95% CrI] |
|---|---|---|---|---|---|---|---|
| Alfalfa | Roza2019 | 424 | 0.61 | 0.95 / 1.25 | 0.26 [0.12, 0.47] | 0.21 [0.09, 0.40] | 0.22 [0.09, 0.40] |
| Blueberry | UF-call† | 1804 | 0.41 | 0.97 / 1.88 | 0.15 [0.09, 0.25] | 0.13 [0.07, 0.21] | 0.12 [0.06, 0.19] |
| Blueberry | UF-fruit | 1559 | 0.42 | 0.93 / 1.34 | 0.17 [0.09, 0.30] | 0.14 [0.07, 0.23] | 0.13 [0.07, 0.22] |
| Blueberry | UF-GP | 1804 | 0.46 | 0.97 / 2.05 | 0.17 [0.10, 0.29] | 0.14 [0.08, 0.22] | 0.10 [0.06, 0.16] |
| *Miscanthus* | Biomass | 197 | 0.47 | 0.99 / 1.14 | 0.19 [0.09, 0.35] | 0.17 [0.08, 0.31] | 0.16 [0.08, 0.30] |
| *Panicum* | Forage | 530 | 0.50 | 0.95 / 1.54 | 0.22 [0.10, 0.39] | 0.20 [0.09, 0.35] | 0.20 [0.09, 0.35] |
| Potato | SLU | 669 | 0.53 | 0.96 / 2.02 | 0.23 [0.10, 0.40] | 0.19 [0.09, 0.34] | 0.18 [0.09, 0.31] |
| Potato | SolCAP | 187 | 0.49 | 0.90 / 1.96 | 0.21 [0.08, 0.41] | 0.18 [0.08, 0.36] | 0.19 [0.08, 0.37] |
| Rose | Rooting | 106 | 0.50 | 0.93 / 1.58 | 0.24 [0.09, 0.48] | 0.22 [0.08, 0.43] | 0.21 [0.08, 0.42] |

†Same individuals as UF-GP, processed with a different dosage-calling pipeline.

##### C.2 Structure-control sensitivity

Figure S1 shows the fitted standard $D$-kernel share under no control, ten genomic principal components and, where available, family, cohort or group-kernel control, for the nine tetraploid diagnostic panels using full-length posterior fits. The $S_{\text{struct}}$ values used for workflow classification are computed separately, from the workflow's own point-estimation refits under the same control specifications, for all eleven panels. Dataset-level values of $S_{\text{struct}}$ and the corresponding structure-ruler thresholds are reported in Table S6.

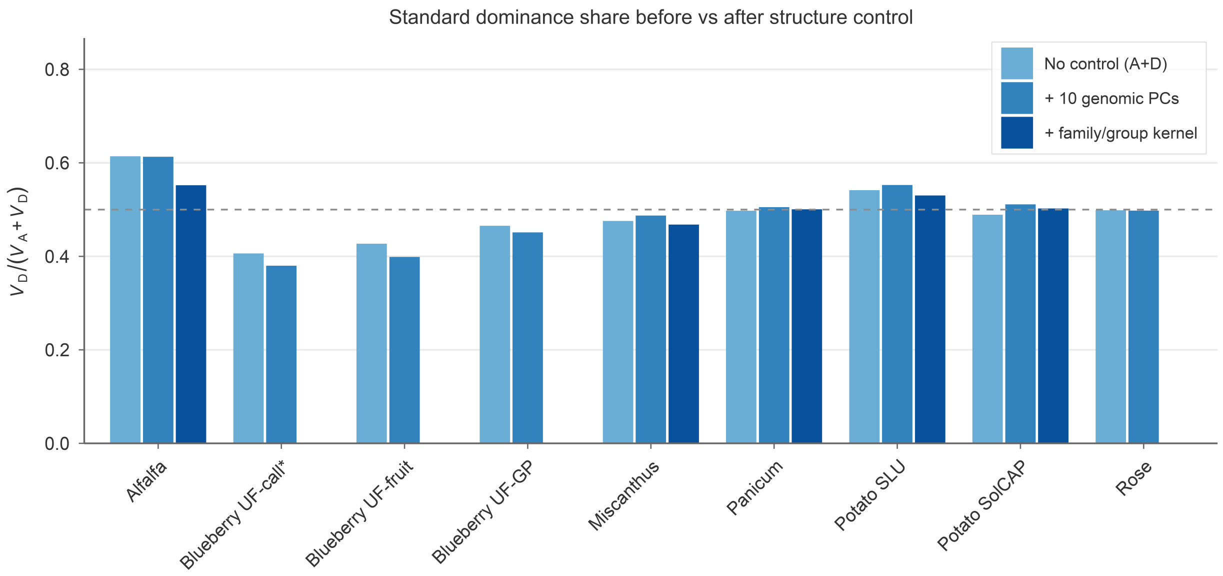


**Figure S1: Standard dominance share under structure control.** For each of the nine tetraploid diagnostic panels, $V_D/(V_A + V_D)$ is shown under no control, with ten genomic principal components and, where available, with a family/group kernel. The dashed line marks 0.5.

##### C.3 Estimator and prior sensitivity of the fitted fractions

Table S5 compares the default-prior Bayesian fractions with fits under changed priors and with REML (methods in Section A.4). Within the Bayesian model, the standard share of a trait differed by at most 0.02 (standard deviation) between seeds and by at most 0.13 between the default and any changed prior, and the orthogonal $D_2$ fraction by at most 0.06. REML intervals for the standard share were wide: of 33 empirical traits, 12 intervals covered the whole range from zero to one, 10 excluded zero (three alfalfa, three potato SLU, two potato SolCAP and two blueberry UF-GP traits) and 13 excluded one. On permuted phenotypes the REML share was at zero or one in 96% of permutations (standard deviation 0.31 to 0.51 per trait), whereas the Bayesian permutation shares had means of 0.49 to 0.54 with standard deviations of 0.015 to 0.034. For the orthogonal model, the REML fraction of degrees three and above was below 0.05 in 26 of 33 traits (three of these round to 0.05 in Table S5), and the REML $D_2$ fraction was 0.49 to 0.55 for the three alfalfa traits and 0.19 to 0.33 for potato SLU, and at most 0.04 in *Panicum*, *Miscanthus*, rose, sweetpotato and sugarcane.

**Table S5: Estimator and prior sensitivity of the fitted variance fractions.** For the first three analysed traits of each panel (one trait for the simulated positive control). *Standard model:* the default-prior `BGLR` share $\widehat{s}_D$ (mean over traits and three seeds); the range of the trait-mean share over four changed prior settings ($R^2 = 0.2$ or 0.8; $\mathrm{df}_0 = 1$ or 30); and the REML share of each trait with its 95% likelihood-ratio interval. *Orthogonal model:* the default-prior `BGLR` fractions of $D_2$, $D_3$ and $D_4$; the range of the trait-mean $D_2$ fraction over the changed priors; and, for each trait, the REML $D_2$ fraction and the summed REML fraction of all higher degrees ($D_2$ / higher). Fractions are of fitted genetic variance. For sugarcane and sweetpotato the orthogonal model has six degrees; degrees five and six are omitted from the `BGLR` column and included in "higher". REML degree fractions are point estimates without intervals. Traits are in the order of the released data files; trait-level values are in the released output.

| | Standard model: share $\widehat{s}_D$ | | | Orthogonal model | | |
|---|---|---|---|---|---|---|
| Panel | `BGLR` | Priors | REML [95% interval], per trait | `BGLR` $D_2$ / $D_3$ / $D_4$ | $D_2$, priors | REML $D_2$ / higher, per trait |
| Alfalfa | 0.58 | 0.54–0.62 | 1.00 [0.50, 1.00]; 1.00 [0.02, 1.00]; 1.00 [0.50, 1.00] | 0.27 / 0.21 / 0.22 | 0.25–0.28 | 0.55 / 0.06; 0.49 / 0.21; 0.54 / 0.07 |
| Blueberry (UF-call†) | 0.41 | 0.33–0.46 | 0.24 [0.00, 0.58]; 0.30 [0.00, 0.68]; 0.12 [0.00, 0.56] | 0.15 / 0.13 / 0.12 | 0.11–0.18 | 0.12 / 0.01; 0.14 / 0.04; 0.05 / 0.23 |
| Blueberry (UF-fruit) | 0.32 | 0.21–0.42 | 0.06 [0.00, 0.42]; 0.00 [0.00, 0.34]; 0.00 [0.00, 0.36] | 0.13 / 0.10 / 0.10 | 0.09–0.18 | 0.09 / 0.00; 0.00 / 0.00; 0.00 / 0.00 |
| Blueberry (UF-GP) | 0.45 | 0.43–0.49 | 0.16 [0.00, 0.50]; 0.44 [0.08, 0.80]; 0.68 [0.14, 1.00] | 0.17 / 0.14 / 0.10 | 0.13–0.20 | 0.12 / 0.05; 0.17 / 0.00; 0.19 / 0.08 |
| *Miscanthus* | 0.47 | 0.43–0.49 | 0.00 [0.00, 1.00]; 0.00 [0.00, 0.72]; 0.00 [0.00, 0.60] | 0.18 / 0.17 / 0.16 | 0.13–0.22 | 0.00 / 0.00; 0.00 / 0.00; 0.00 / 0.00 |
| *Panicum* | 0.49 | 0.48–0.50 | 0.64 [0.00, 1.00]; 0.00 [0.00, 1.00]; 0.02 [0.00, 1.00] | 0.20 / 0.19 / 0.19 | 0.16–0.23 | 0.00 / 0.00; 0.00 / 0.00; 0.00 / 0.00 |
| Potato (SLU) | 0.56 | 0.53–0.59 | 0.72 [0.40, 1.00]; 0.76 [0.40, 1.00]; 0.70 [0.16, 1.00] | 0.21 / 0.16 / 0.16 | 0.19–0.23 | 0.33 / 0.05; 0.29 / 0.22; 0.19 / 0.00 |
| Potato (SolCAP) | 0.50 | 0.47–0.51 | 0.46 [0.00, 1.00]; 0.40 [0.10, 0.74]; 0.88 [0.40, 1.00] | 0.20 / 0.18 / 0.18 | 0.18–0.23 | 0.72 / 0.00; 0.12 / 0.00; 0.22 / 0.00 |
| Rose | 0.48 | 0.47–0.49 | 0.74 [0.00, 1.00]; 0.00 [0.00, 1.00]; 0.00 [0.00, 1.00] | 0.22 / 0.21 / 0.20 | 0.20–0.24 | 0.04 / 0.00; 0.00 / 0.20; 0.00 / 0.00 |
| Sugarcane | 0.48 | 0.47–0.50 | 0.00 [0.00, 1.00]; 0.48 [0.00, 1.00]; 0.10 [0.00, 1.00] | 0.17 / 0.16 / 0.17 | 0.14–0.18 | 0.00 / 0.00; 0.00 / 0.05; 0.00 / 0.00 |
| Sweetpotato | 0.47 | 0.41–0.49 | 0.80 [0.00, 1.00]; 0.00 [0.00, 0.76]; 0.10 [0.00, 0.94] | 0.10 / 0.09 / 0.09 | 0.07–0.13 | 0.00 / 0.00; 0.00 / 0.00; 0.00 / 0.00 |
| Positive control (sim) | 0.77 | 0.66–0.83 | 1.00 [0.92, 1.00] | 0.73 / 0.09 / 0.08 | 0.66–0.81 | 0.93 / 0.05 |

†Same individuals as UF-GP, processed with a different dosage-calling pipeline.

#### C.4 Full panel-conditioned workflow scorecard

Table S6 reports the principal decision variables used by the panel-conditioned diagnostic workflow; the full scorecard, including the additive-null and spike-in reference distributions behind the derived criteria, is distributed with the released code. Columns are grouped by criterion, in workflow order: kernel geometry, structure sensitivity, standard $D$-kernel share relative to the phenotype-permutation no-signal null, orthogonal $D_2$ localization, planted-$D_2$ spike-in recoverability and paired cross-validated prediction gain. The final column gives the reporting label assigned to each panel. Threshold values used for label assignment are listed in Table S7. Blueberry UF-GP was the only empirical panel that passed the additive-null percentile and the interval-width limit; it failed the permutation-null margin, and its $D_2$ fraction (0.17) was closer to the additive-only median (0.09) than to the spike-in median (0.37), so the localization criterion on $P_{D_2}$ was not met either. The two reference medians are in the released scorecard and are not tabulated here.

For the nine tetraploid panels, $P_{D_2}$, $L_{D_2}$ and $W_{D_2}$ are computed from the trait-averaged degree fractions and interval endpoints, so $L_{D_2}$ is the difference of trait means and not the mean of trait-level differences; for sugarcane and sweetpotato they are means of the trait-level values.

The confidence interval of the prediction gain $\Delta_{A+D}$ is the mean $\pm 1.96$ standard errors over all trait-by-repeat contrasts, which treats repeats of the same trait as replicates; it is narrower than an interval based on between-trait variation in nine of the eleven empirical panels. With a between-trait standard error the lower bound stays above zero for alfalfa (+0.041) and potato SLU (+0.003) but not for potato SolCAP (−0.011) or sugarcane (−0.013), so the *prediction-useful* label of the latter two depends on this choice.

**Table S6: Full panel-conditioned workflow scorecard.** For each empirical dataset and for the simulated positive control, the table reports the decision variables used by the workflow in Figure 6. Columns are grouped by criterion, in workflow order, and the final column gives the resulting workflow label. Column definitions are given below the table.

| | Geometry | | Structure | | Standard share vs. permutation null | | | Orthogonal $D_2$ localization | | | | Recov. | Pred. | |
|---|---|---|---|---|---|---|---|---|---|---|---|---|---|---|
| Panel | A–D cos | $r_{\text{eff}}$ | $S_{\text{struct}}$ | $q_{0.95}^{\text{struct}}$ | $\widehat{s}_{D,\text{real}}$ | Null | $\Delta_{\text{null}}$ | $P_{D_2}$ | $L_{D_2}$ | $q_{L,\text{add0}}$ | $W_{D_2}$ | $R_{D_2}$ | $\Delta_{A+D}$ [95% lo] | Label |
| Alfalfa | 0.95 | 1.25 | 0.01 | 0.12 | 0.59 | 0.53 | +0.058 | 0.26 | +0.045 | 0.87 | 0.35 | 0.83 | +0.052 [+0.042] | prediction-useful |
| Blueberry (UF-call†) | 0.97 | 1.88 | 0.07 | 0.11 | 0.43 | 0.51 | −0.083 | 0.15 | +0.020 | 0.83 | 0.17 | 1.00 | −0.001 [−0.002] | model-basis |
| Blueberry (UF-fruit) | 0.93 | 1.34 | 0.04 | 0.14 | 0.31 | 0.54 | −0.224 | 0.17 | +0.033 | 0.87 | 0.21 | 1.00 | −0.001 [−0.002] | model-basis |
| Blueberry (UF-GP) | 0.97 | 2.05 | 0.03 | 0.07 | 0.46 | 0.51 | −0.056 | 0.17 | +0.039 | 1.00 | 0.19 | 1.00 | +0.003 [+0.001] | model-basis |
| *Miscanthus* | 0.99 | 1.14 | 0.02 | 0.07 | 0.48 | 0.49 | −0.018 | 0.19 | +0.020 | 0.47 | 0.27 | 0.73 | 0.000 [−0.004] | model-basis‡ |
| *Panicum* | 0.95 | 1.54 | 0.02 | 0.12 | 0.49 | 0.50 | −0.011 | 0.22 | +0.018 | 0.73 | 0.29 | 0.90 | +0.002 [−0.001] | model-basis |
| Potato (SLU) | 0.96 | 2.02 | 0.05 | 0.11 | 0.58 | 0.52 | +0.067 | 0.23 | +0.035 | 0.93 | 0.29 | 1.00 | +0.012 [+0.006] | prediction-useful |
| Potato (SolCAP) | 0.90 | 1.96 | 0.00 | 0.10 | 0.48 | 0.52 | −0.046 | 0.21 | +0.025 | 0.73 | 0.32 | 0.73 | +0.020 [+0.007] | prediction-useful‡ |
| Rose | 0.93 | 1.58 | 0.01 | 0.08 | 0.50 | 0.52 | −0.018 | 0.24 | +0.027 | 0.87 | 0.39 | 0.67 | −0.007 [−0.020] | model-basis‡ |
| Sugarcane | 0.97 | 1.85 | 0.00 | 0.05 | 0.48 | 0.49 | −0.016 | 0.18 | +0.007 | 0.30 | 0.37 | 0.77 | +0.011 [+0.0002] | prediction-useful‡ |
| Sweetpotato | 0.96 | 1.23 | 0.00 | 0.06 | 0.48 | 0.50 | −0.027 | 0.11 | +0.013 | 0.27 | 0.17 | 0.87 | +0.002 [−0.0004] | model-basis |
| Positive control (sim)§ | 0.77 | 1.84 | 0.00 | 0.04 | 0.78 | 0.51 | +0.270 | 0.74 | +0.647 | 1.00 | 0.14 | 1.00 | +0.191 [+0.179] | $D_2$-supported |

Values are dataset-level means across analysed traits where applicable (alfalfa: its representative yield trait for the localization columns). $\widehat{s}_{D,\text{real}}$ and "Null" are the observed and the mean permuted standard $D$-kernel share on the first three analysed traits, computed with the permutation point estimator, so $\widehat{s}_{D,\text{real}}$ can differ from the all-trait share in Table S4. The bracketed value is the lower bound of the 95% confidence interval of $\Delta_{A+D}$. Other workflow quantities are defined in Section A.5 and their thresholds are listed in Table S7. †Same individuals as UF-GP, processed with a different dosage-calling pipeline. ‡Flagged *unresolved by panel geometry* (a planted $D_2$ spike-in is not recoverable). §Simulated panel with one trait and no family grouping (Section A.2).

**Table S7:** Threshold values used in the panel-conditioned diagnostic workflow.

| **Diagnostic quantity** | **Threshold** | **Use in workflow** |
|---|---|---|
| Structure sensitivity, $S_{\text{struct}}$ | $\geq 0.10$ or $> q_{0.95}^{\text{struct}}$ | Flags fitted dominance share as structure-sensitive |
| Permutation-null excess, $\Delta_{\text{null}}$ | $\Delta_{\text{null}} > 0.05$ | Required for $D_2$-supported interpretation |
| Spike-in recovery, $R_{D_2}$ | $\geq 0.80$ | Required for $D_2$-supported interpretation |
| $D_2$ localization, $P_{D_2}$ | Closer to planted-$D_2$ spike-in median than to additive-only null median | Required for $D_2$-supported interpretation |
| Orthogonal $D_2$ excess percentile, $q_{L,\text{add0}}$ | $> 0.95$ | Required for $D_2$-supported interpretation |
| Posterior uncertainty, $W_{D_2}$ | $< 0.25$ | Required for $D_2$-supported interpretation |
| Prediction gain, $\Delta_{A+D}$ | $\geq 0.01$ with lower confidence bound $> 0$ | Classifies non-$D_2$-supported estimates as prediction-useful |

#### C.5 Sensitivity of the labels to the thresholds

The thresholds in Table S7 are conventions, so we re-applied the label rule to the scorecard with each of the six numerical thresholds lowered and raised by 20%: the absolute structure margin, the permutation-null margin, the spike-in recovery rate, the additive-null percentile, the credible-interval width and the prediction gain. The additive-null percentile cannot exceed one, so its raised value was set to 0.999 (above all 30 control replicates) and not to 1.14. No model was refitted. Changing one threshold at a time altered two of 132 panel-by-scenario combinations for the eleven empirical panels: sugarcane (gain 0.0113) and potato SLU (gain 0.01198) moved from *prediction-useful* to *model-basis* when the prediction-gain threshold was raised to 0.012. Changing all six thresholds together, over the 64 combinations of lowered and raised values, left nine panels with the same label in every combination. Sugarcane was *model-basis* in the 32 combinations with the raised prediction-gain threshold. Potato SLU was labelled $D_2$*-supported* in the 16 combinations that both widened the credible-interval limit to 0.30 and lowered the additive-null percentile to 0.76, and *model-basis* in 24 of the remaining 48. It is the empirical panel closest to $D_2$ support, failing the default rule on credible-interval width (0.295 against 0.25) and on the additive-null percentile (0.93 against 0.95). The labels of the other panels, including the absence of any *structure-sensitive* label, did not depend on the thresholds within this range. The simulated positive control kept its $D_2$*-supported* label in every combination.

### Supplemental references


Amadeu, R. R., Garcia, A. A. F., Munoz, P. R., & Ferrão, L. F. V. (2023). AGHmatrix: genetic relationship matrices in R. *Bioinformatics*, *39*(7), btad445. https://doi.org/10.1093/bioinformatics/btad445

Batista, L. G., Mello, V. H., Souza, A. P., & Margarido, G. R. A. (2022). Genomic prediction with allele dosage information in highly polyploid species. *Theoretical and Applied Genetics*, *135*(2), 723–739. https://doi.org/10.1007/s00122-021-03994-w

de Bem Oliveira, I., Amadeu, R. R., Ferrão, L. F. V., & Muñoz, P. R. (2020). Optimizing whole-genomic prediction for autotetraploid blueberry breeding. *Heredity*, *125*(6), 437–448. https://doi.org/10.1038/s41437-020-00357-x

Endelman, J. B., Schmitz Carley, C. A., Bethke, P. C., Coombs, J. J., Clough, M. E., da Silva, W. L., De Jong, W. S., Douches, D. S., Frederick, C. M., Haynes, K. G., Holm, D. G., Miller, J. C., Muñoz, P. R., Navarro, F. M., Novy, R. G., Palta, J. P., Porter, G. A., Rak, K. T., Sathuvalli, V. R., . . . Yencho, G. C. (2018). Genetic variance partitioning and genome-wide prediction with allele dosage information in autotetraploid potato. *Genetics*, *209*(1), 77–87. https://doi.org/10.1534/genetics.118.300685

Ferrão, L. F. V., Benevenuto, J., de Bem Oliveira, I., Cellon, C., Olmstead, J., Kirst, M., Resende, M. F. R., Jr., & Muñoz, P. (2018). Insights into the genetic basis of blueberry fruit-related traits using diploid and polyploid models in a GWAS context. *Frontiers in Ecology and Evolution*, *6*, 107. https://doi.org/10.3389/fevo.2018.00107

Gaynor, R. C., Gorjanc, G., & Hickey, J. M. (2021). AlphaSimR: an R package for breeding program simulations. *G3: Genes, Genomes, Genetics*, *11*(2), jkaa017. https://doi.org/10.1093/g3journal/jkaa017

Gerard, D., Ferrão, L. F. V., Garcia, A. A. F., & Stephens, M. (2018). Genotyping polyploids from messy sequencing data. *Genetics*, *210*(3), 789–807. https://doi.org/10.1534/genetics.118.301468

Lara, L. A. d. C., Santos, M. F., Jank, L., Chiari, L., Vilela, M. d. M., Amadeu, R. R., dos Santos, J. P. R., Pereira, G. d. S., Zeng, Z.-B., & Garcia, A. A. F. (2019). Genomic selection with allele dosage in Panicum maximum Jacq. *G3: Genes, Genomes, Genetics*, *9*(8), 2463–2475. https://doi.org/10.1534/g3.118.200986

Medina, C. A., Hansen, J., Crawford, J., Viands, D., Sapkota, M., Xu, Z., Peel, M. D., & Yu, L.-X. (2025). Genome-wide association and genomic prediction of alfalfa (Medicago sativa L.) biomass yield under drought stress. *International Journal of Molecular Sciences*, *26*(2), 608. https://doi.org/10.3390/ijms26020608

Njuguna, J. N., Clark, L. V., Lipka, A. E., Anzoua, K. G., Bagmet, L., Chebukin, P., Dwiyanti, M. S., Dzyubenko, E., Dzyubenko, N., Ghimire, B. K., Jin, X., Johnson, D. A., Kjeldsen, J. B., Nagano, H., de Bem Oliveira, I., Peng, J., Petersen, K. K., Sabitov, A., Seong, E. S., ... Sacks, E. J. (2023). Impact of genotype-calling methodologies on genome-wide association and genomic prediction in polyploids. *The Plant Genome*, *16*(4), e20401. https://doi.org/10.1002/tpg2.20401

Pérez, P., & de los Campos, G. (2014). Genome-wide regression and prediction with the BGLR statistical package. *Genetics*, *198*(2), 483–495. https://doi.org/10.1534/genetics.114.164442

Rosyara, U. R., De Jong, W. S., Douches, D. S., & Endelman, J. B. (2016). Software for genome-wide association studies in autopolyploids and its application to potato. *The Plant Genome*, *9*(2), plantgenome2015.08.0073. https://doi.org/10.3835/plantgenome2015.08.0073

Selga, C., Reslow, F., Pérez-Rodríguez, P., & Ortiz, R. (2022). The power of genomic estimated breeding values for selection when using a finite population size in genetic improvement of tetraploid potato. *G3: Genes, Genomes, Genetics*, *12*(1), jkab362. (Data: figshare `10.25387/g3.14376938`) https://doi.org/10.1093/g3journal/jkab362

Slater, A. T., Cogan, N. O. I., Forster, J. W., Hayes, B. J., & Daetwyler, H. D. (2016). Improving genetic gain with genomic selection in autotetraploid potato. *The Plant Genome*, *9*(3), plantgenome2016.02.0021. https://doi.org/10.3835/plantgenome2016.02.0021

Wamhoff, D., Marxen, A., Acharya, B., Grzelak, M., Debener, T., & Winkelmann, T. (2024). *Genotypic and phenotypic raw data for GWAS study Wamhoff et al. 2024.* (Research Data Repository, Leibniz Universität Hannover (LUIS)) https://doi.org/10.25835/xvdntbqg
